# Structure-guided function discovery of an NRPS-like glycine betaine reductase for choline biosynthesis in fungi

**DOI:** 10.1101/559534

**Authors:** Yang Hai, Arthur Huang, Yi Tang

## Abstract

Nonribosomal peptide synthetases (NRPS) and NRPS-like enzymes have diverse functions in primary and secondary metabolism. By using a structure-guided approach, we uncovered the function of an NRPS-like enzyme with unusual domain architecture, catalyzing two sequential two-electron reductions of glycine betaine to choline. Structural analysis based on homology model suggests cation-π interactions as the major substrate specificity determinant, which was verified using substrate analogs and inhibitors. Bioinformatic analysis indicates this NRPS-like glycine betaine reductase is highly conserved and widespread in fungi kingdom. Genetic knockout experiments confirmed its role in choline biosynthesis and maintaining glycine betaine homeostasis in fungi. Our findings demonstrate that the oxidative choline-glycine betaine degradation pathway can operate in a fully reversible fashion and provide new insights in understanding fungal choline metabolism. The use of an NRPS-like enzyme for reductive choline formation is energetically efficient compared to known pathways. Our discovery also underscores the capabilities of structure-guided approach in assigning function of uncharacterized multidomain proteins, which can potentially aid functional discovery of new enzymes by genome mining.

## Introduction

The nonribosomal peptide synthetase (NRPS)-like carboxylic acid reductases (CARs) catalyze ATP-and NADPH-dependent reduction of carboxylic acids to the corresponding aldehydes.^1^ These multi-domain enzymes consist of an *N*-terminal adenylation domain (A), which selects and activates carboxylic acid substrate by adenylation, then transfers the acyl moiety to the phosphopantetheinyl arm from the thiolation domain (T) through a thioester linkage. The T domain delivers the acyl thioester to the *C*-terminal thioester reductase domain (R), where the NADPH-dependent thioester reduction occurs (Figure 1A). The R domains in CARs catalyze a strict two-electron reduction without over-reducing the aldehyde product to alcohol and the structural basis for such specificity was reported recently.^2^

**Figure 1.**
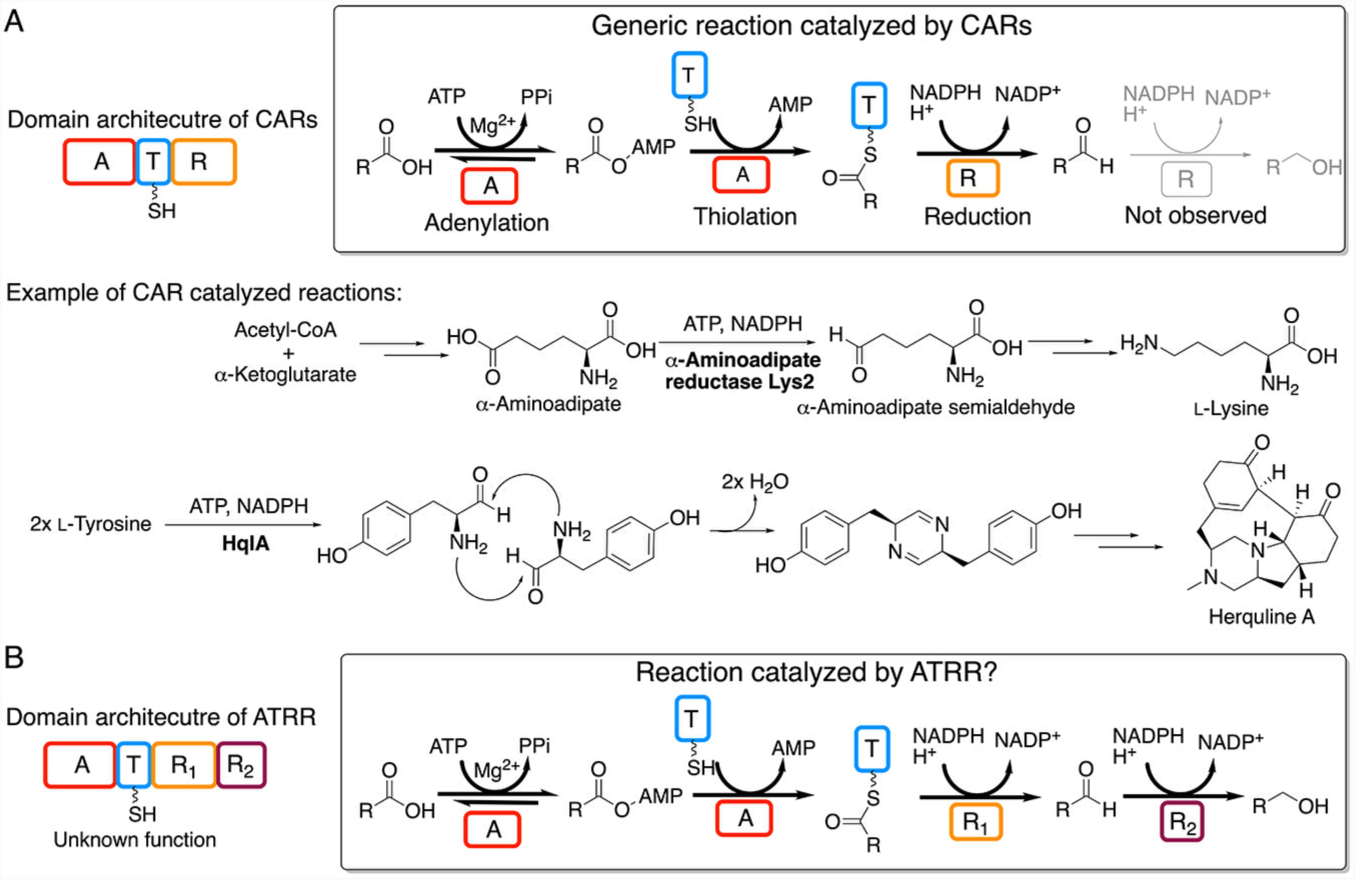
Comparison between typical CARs and ATRR. (*A*) Domain architecture of typical CARs and CAR-catalyzed reactions. A, adenylation domain; T, thiolation domain, R, thioester reductase domain. (*B*) Domain architecture of ATRR and the postulated reaction based on domain function.

The members of the CAR family are functionally diverse and play indispensable roles in both primary and secondary metabolism, such as the α-aminoadipate reductase Lys2 required for lysine biosynthesis in fungi;^3^ and tyrosine reductase HqlA involved in biosynthesis of herquline A (Figure 1A).^4^ CARs also show substantial promise as green biocatalysts for the conversion of aromatic, or aliphatic carboxylic acids into the corresponding aldehydes. Indeed, synthetic applications of CARs have been reported for biocatalytic synthesis of vanillin;^5^ preparation of methyl branched aliphatic aldehydes;^6^ production of biofuels,^7-8^ and other chemical commodities.^9^

Prominent metabolic roles of CARs and their biocatalytic potential encouraged us to discover new members with novel functions by genome mining. During this effort, we focused on a fungal CAR-like protein with unknown function. It is distinguished from all other CARs due to an extra *C*-terminal YdfG-like short-chain dehydrogenase/reductase domain. We named this protein ATRR and the corresponding gene *atrr* after its unusual domain architecture (A-T-R_1_-R_2_). The presence of two fused R domains in ATRR implies that it could catalyze two consecutive two-electron reductions of a carboxylic acid to yield an alcohol (Figure 1B). Moreover, genomic survey reveals that *atrr* genes are widespread but exclusive to eukaryotes, mostly in fungi and also found in several protists and invertebrates species. In particular, ATRR orthologues are highly conserved in the fungi kingdom: sequence identities are greater than 60% from different species, which indicates a unified and conserved role of ATRR. Transcriptomics studies also suggest *atrr* genes are constitutively transcribed in different fungal species, including human opportunistic pathogen *Aspergillus fumigatus*; ^10,11^ and rice blast fungus *Magnaporthe oryzae*.^12^ These features motivated us to discover and characterize its enzymatic activity and biological function.

## RESULTS

### Bioinformatic analysis provides initial insight

We first applied the “genomic enzymology” strategy to obtain clues regarding ATRR function from *atrr* genomic context, since enzymes in a microbial metabolic pathway often are encoded by a gene cluster.^13,14^ However, comparative genomic analysis of *atrr* orthologues in fungi reveals that there are no well-defined and conserved gene neighborhoods. Even in some close-related species of the same genus (e.g. *Aspergillus*), the genome environment of *atrr* differs substantially (*SI Appendix*, Fig. S2-3). The diversified genomic context and lack of obvious co-occurrence with signature metabolic pathway genes prevented us from reliable functional assignment.

We then focused on characterization of the A domain of ATRR (ATRR-A), which selects substrate and is therefore the entry point to ATRR catalysis. Because the substrate specificity of microbial A domain is dictated by the amino acid residues lining the active site pocket (10 AA code), deciphering this “nonribosomal code” of ATRR-A would in theory allow us to predict its function.^15,16^ Bioinformatic analysis showed the ATRR-A has a unique 10AA code, and it is conserved among ATRR orthologues from different fungal species (*SI Appendix*, Fig. S4). However, substrate prediction by online servers was unsuccessful due to the lack of characterized homologues with significant similarity.^17,18^ Closer inspection reveals that the highly conserved aspartate residue (D235 in PheA) which typically interacts with substrate α-amino group via a salt bridge,^19,20^ is not conserved in in ATRR-A (Figure 2A). Loss of this key aspartate residue suggests that the substrate is not a typical α-amino acid, or an unusual amino acid substrate accommodated in an unprecedented fashion, as in the case of ClbH-A2 which activates *S*-adenosyl-L-methionine (SAM).^21^ Indeed, most characterized A domains without D235 equivalent residues (Asp/Glu) are known to activate non-amino acid type substrates, including aryl acid, keto acid, and hydroxy acid (Figure 2A).^22-25^ Based on this prediction, we screened the CAR activity of ATRR *in vitro* against a library of common carboxylic acids from primary metabolism, as well as the twenty proteinogenic amino acids (*SI Appendix*, Fig. S5). The *A. nidulans* ATRR protein (Uniprot AN5318.2) was expressed from *E. coli* BL21(DE3) and purified to homogeneity by affinity chromatography and size-exclusion chromatography (*SI Appendix*, Fig. S6). *Apo*-ATRR was enzymatically converted to *holo*-form by using phosphopantetheinyl transferase NpgA and coenzyme A (*SI Appendix*, Fig. S7).^26^ The reductase activity of *holo*-ATRR was examined by monitoring the oxidation of NADPH at 340 nm. ATRR showed no activity towards any of the substrates in this library.

**Figure 2.**
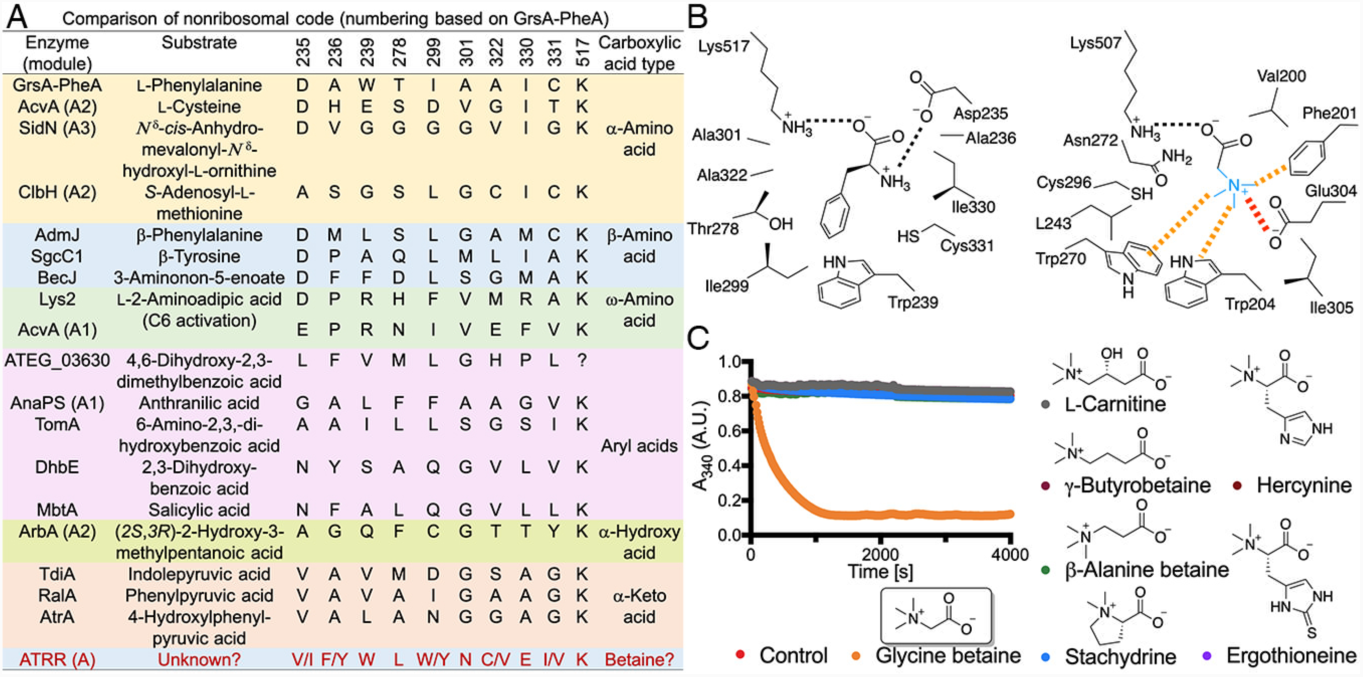
Structure-based prediction of ATRR substrate. (*A*) Summary of the 10-AA code of ATRR-A and selected adenylation domains. The K517 equivalent residue in ATEG_03630 was unidentified. (*B*) Comparison of the homology model of *A. nidulans* ATRR-A domain with structure of PheA in complex with its cognate substrate L-phenylalanine. The conserved salt bridges recognizing the α-carboxylate and α-amino group are shown as black dashes. The proposed cation-π and electrostatic interactions favoring binding of betaine-like substrates in ATRR A domain are shown as orange and red dashes, respectively. The simplest betaine, glycine betaine, was shown to indicate the plausible binding mode. (*C*) Carboxylic acid reductase activity of ATRR against a panel of betaine substrates.

### Structure-based approach predicts A domain substrate

We then used a structure-based approach to gain more insights about the substrate specificity of ATRR-A, a strategy that has led to many successful predictions for enzymes of unknown function.^27,28^ We constructed a homology model of ATRR-A by using a TycA variant (PDB entry 5N82)^25^ as the template (34% sequence identity to ATRR-A), followed by manual replacement of the 10 AA code residues to the corresponding ones in *A. nidulans* ATRR-A (Figure 2B). In comparison to the structure of the archetypical adenylation domain PheA,^15^ three aromatic residues (F201, W204, W270) stood out in the ATRR-A homology model.

These residues could potentially form an aromatic cage to bind a quaternary ammonium group through cation-π interactions.^29,30^ This quaternary ammonium group can be further electrostatically stabilized by a nearby anionic residue E304. Such aromatic box is reminiscent of those observed in quaternary ammonium moiety binding proteins, such as periplasmic betaine binding protein Prox;^31^ *cis*-4-hydroxy-D-proline betaine epimerase HpbD, and demethylase HpbJ;^27^ histone trimethyllysine reader proteins;^32^ and acetylcholine esterase.^29,30^

We therefore speculated that the carboxylic acid substrate of ATRR could be a small betaine, such as the physiologically abundant carnitine, glycine betaine, etc. To test this hypothesis, we assayed ATRR activity with a panel of naturally occurring betaines (Figure 2C). Gratifyingly, rapid oxidation of NADPH was observed only in the presence of the smallest betaine, glycine betaine. Consumption of NADPH indicates that glycine betaine was reduced. In accordance with our early hypothesis about ATRR function (Figure 1B), the predicted corresponding [2+2]-electron reduction product choline was indeed accumulated in the reaction mixture (*SI Appendix*, Fig. S8). We then determined the steady-state kinetic constants for ATRR glycine betaine reductase activity (Table 1, *SI Appendix*, Fig. S9). The overall *k*_cat_ value (∼1 s^−1^) is modest but similar to that of previously characterized CARs,^2^ while the *K*_m_ value (∼2 mM) matches the intracellular level of glycine betaine (∼ 3 mM) in fungi.^33^ Therefore, glycine betaine is a physiologically relevant substrate for ATRR.

**Table 1.** Steady-state kinetic parameters for ATRR with different substrates.

| Enzyme construct | Substrate | $k_{\text{cat}}$ ($\text{s}^{-1}$) | $K_{\text{M}}$ (mM) | $k_{\text{cat}}/K_{\text{M}}$ ($\text{M}^{-1}\text{s}^{-1}$) |
| --- | --- | --- | --- | --- |
| ATR<sub>1</sub>R<sub>2</sub> | Glycine betaine | $0.94 \pm 0.03$ | $2.1 \pm 0.2$ | $4.5 \times 10^2$ |
| | Glycine betaine aldehyde | $7.1 \pm 0.3$ | $0.24 \pm 0.03$ | $3.0 \times 10^4$ |
| | *N,N*-Dimethylglycine<sup>1</sup> | $0.023 \pm 0.002$ | $5.8 \pm 0.8$ | 4.0 |
|  | 3,3,-Dimethylbutyrate | inactive | inactive | inactive |
| ATR<sub>1</sub><sup>0</sup>R<sub>2</sub> (Y811F) | Glycine betaine | inactive | inactive | inactive |
| | Glycine betaine aldehyde | $7.0 \pm 0.2$ | $0.24 \pm 0.03$ | $2.9 \times 10^4$ |
| ATR<sub>1</sub>R<sub>2</sub><sup>0</sup> (Y1178F) | Glycine betaine | $0.72 \pm 0.02$ | $2.2 \pm 0.2$ | $3.3 \times 10^2$ |
| | Glycine betaine aldehyde | $0.30 \pm 0.04$ | $1.5 \pm 0.6$ | $1.9 \times 10^2$ |
| ATR<sub>1</sub>R<sub>2</sub><sup>0</sup> (G1036A/Y1178F) | Glycine betaine | $0.66 \pm 0.02$ | $2.2 \pm 0.3$ | $3.0 \times 10^2$ |
| | Glycine betaine aldehyde | $0.018 \pm 0.001$ | $2.6 \pm 0.5$ | 7.0 |
| R<sub>2</sub> | Glycine betaine | inactive | inactive | inactive |
| | Glycine betaine aldehyde | $7.5 \pm 0.2$ | $0.28 \pm 0.03$ | $2.7 \times 10^4$ |
| | 3,3,-Dimethylbutyraldehyde | $0.20 \pm 0.02$ | $90 \pm 20$ | 2 |

<sup>1</sup>Substrate inhibition ( $K_{\text{i}} = 48 \pm 9$ mM) was observed.

### Mechanistic study of ATRR

We next studied the catalytic mechanism of the glycine betaine reductase function of ATRR. By using a hydroxylamine-trapping assay designed for testing adenylation activity,^34^ we confirmed that glycine betaine was indeed adenylated by ATRR-A (Figure 3A). Furthermore, similar to all CARs, the glycine betaine reductase activity of ATRR requires phosphopantetheinylation of the T domain, suggesting the corresponding glycine betainoyl thioester intermediate is critical for reduction (*SI Appendix*, Fig. S10). We verified the formation of this intermediate using MALDI-TOF mass spectrometry with the standalone ATRR-T domain, which can be acylated by ATRR-A in the presence of glycine betaine and ATP (*SI Appendix*, Fig. S7). The expected on-pathway intermediate, glycine betaine aldehyde (hydrate), was also observed during the reaction course by continuous trapping with phenylhydrazine (*SI Appendix*, Fig. S11). Direct reduction of glycine betaine aldehyde to choline by ATRR was also observed, confirming its role as an intermediate in the step-wise reduction (*SI Appendix*, Fig. S8). Kinetic assay showed that glycine betaine aldehyde reduction by ATRR is more efficient: the catalytic efficiency (*k*_cat_/*K*_m_ = 3.0 × 10^4^ M^−1^s^−1^) is 66-fold greater than that of the overall carboxylic acid reduction while the turnover number (*k*_cat_ = 7 s^−1^) is 6-fold higher. Rapid reduction of glycine betaine aldehyde therefore shows the aldehyde reduction step is not rate-limiting, thereby avoiding accumulation of reactive aldehyde species.

**Figure 3.**
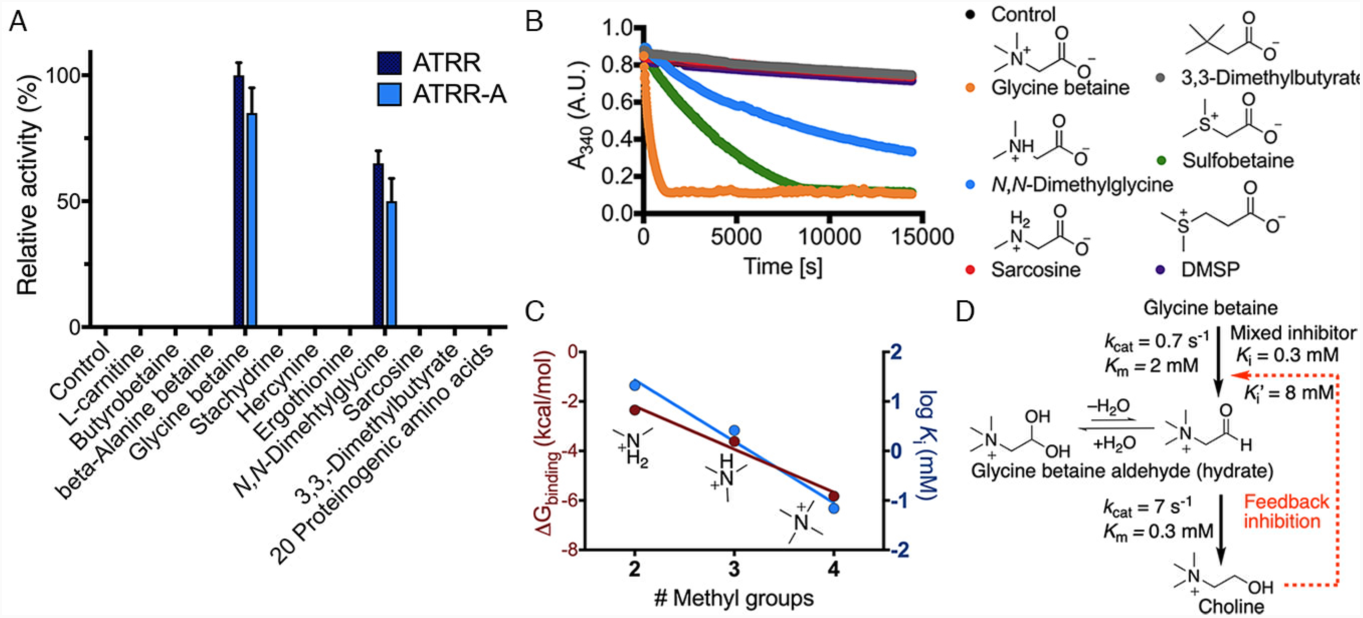
ATRR substrate specificity and inhibition studies. (*A*) Substrate specificity of ATRR-A profiled with different substrates. (*B*) Carboxylic acid reductase activity of ATRR against a panel of glycine betaine analogues. (*C*) Dependence of binding free energy change and inhibitor disassociation constant *K*_*i*_ on the number of methyl groups of ammonium inhibitors against glycine betaine reductase activity. Note that for mixed inhibitor trimethylammonium ion *K*_i_’ is not taken into account. (D) Feedback inhibition from choline could regulate ATRR enzyme activity.

We next characterized the individual function of R_1_ and R_2_ domains. In order to isolate the catalytic activity of individual domain, we inactivated each reductase domain by substituting the conserved catalytic tyrosine residue with phenylalanine (Y811F in R_1_ and Y1178F in R_2_). In agreement with our two sequential two-electron reductions hypothesis (Figure 1B), mutation of R_1_ essentially abolished overall carboxylic acid reductase activity, whereas aldehyde reductase activity remained intact (Table 1). On the contrary, mutation of R_2_ did not substantially affect carboxylic acid reductase activity but resulted in accumulation of glycine betaine aldehyde intermediate (*SI Appendix*, Fig. S11). The Y1178F mutation in R_2_ also led to greater than 150-fold loss of aldehyde reductase activity (catalytic efficiency). (Table 1). To further confirm that the remaining aldehyde reductase activity was not contributed from R_1_, we introduced another mutation at R_2_ (G1036A) to compromise binding of the co-substrate NADPH. As expected, double mutation (G1036A/Y1178F) in R_2_ further decreased the aldehyde reductase activity by 4000-fold, while the carboxylic acid reductase activity was not compromised. The role of R_2_ as a dedicated aldehyde reductase was also supported by the robust reduction of betaine aldehyde to choline using the excised, stand-alone R_2_ domain (Table 1). In summary, the thioester reductase domain R_1_ strictly carries out a single two-electron reduction of glycine betainoyl thioester to glycine betaine aldehyde, which is then reduced to choline at the aldehyde reductase domain R_2_.

### Substrate specificity and inhibition of ATRR

Establishing the function of ATRR as glycine betaine reductase enabled us to further characterize its substrate specificity. We first studied ATRR specificity towards glycine betaine by assaying both adenylation and carboxylic acid reductase activity of ATRR against different glycine betaine analogues (Figure 3A, B). The isosteric, neutral analogue 3,3-dimethylbutyrate completely abolished the activity. In contrast, the tertiary sulfonium analogue sulfobetaine, which also bears a positive charge on its head group but lacks one methyl group compared to glycine betaine, was partially active (an order of magnitude slower). Such comparison clearly demonstrates that the formal positive charge carried by substrate is critical for activity, and thus supports our predicted cation-π interactions conferred by ATRR-A during substrate recognition. The decreased activity of ATRR towards sulfobetaine suggests the importance of the trimethyl moiety for function. The important contribution of the trimethyl moiety of glycine betaine to ATRR catalysis was also revealed by carboxylic acid reduction assays with dimethylglycine and monomethylglycine (sarcosine), which are physiologically relevant analogues as they are intermediates in the glycine betaine degradation pathway.^35^ Progressive decreasing the number of *N-*methyl groups of glycine betaine resulted in significant loss of carboxylic acid reductase activity: the reduction of dimethylglycine was two orders of magnitude slower than that of glycine betaine, while no reduction could be observed with sarcosine (Figure 3A, B, Table 1). Since the ammonium groups of both dimethylglycine (pKa = 9.89) and sarcosine (pKa = 10.1) were thought to be protonated under our experimental condition (pH 7.5 – 8.0),^36^ the decrease of activity per methyl group reflects the additive contribution of each methyl substitution to catalysis. ATRR-A was also shown to be strictly size-selective in that β-alanine betaine and dimethylsulfoniopropionate (DMSP), each containing one additional methylene group to glycine betaine and sulfobetaine, respectively, were not substrates of ATRR. Collectively, these results show that ATRR is highly specific to glycine betaine and such specificity is likely achieved by favoring binding of its head-group and distinguishing different extent of methylation.

To corroborate our findings on substrate specificity, we studied the inhibition of ATRR by using different ammonium-containing inhibitors (Table 2, *SI Appendix*, Fig. S12). Tetramethylammonium ion, which mimics the head-group of glycine betaine, was found to be a potent competitive inhibitor of ATRR-A (*K*_i_ = 69 µM), whereas the potency decreased by 3 orders of magnitude with dimethylammonium (*K*_i_ = 21 mM). As illustrated in Figure 3C, monotonic increase in inhibitor affinity is correlated with addition of each methyl group yielding an incremental free energy relationship: addition of each methyl group contributes approximately 1.7 kcal/mol towards inhibitor binding. Therefore, our results reinforced the idea that the head-group plays a major role in ATRR-A substrate recognition and ATRR-A favors binding of tetraalkylated quaternary ammonium moiety, presumably through a combination of cation-π interactions, hydrophobic effect, and maximized van der Waals interactions. Unlike ATRR-A, R_2_ is not potently inhibited by the head-group mimic tetramethylammonium ion (mixed inhibitor, *K*_i_ = 5 mM, *K*_i_ ^’^ = 18 mM), suggesting the head-group is not the dominant determinant for R_2_ substrate specificity. Nonetheless, the positive charge on the substrate is still essential for R_2_ catalysis as the neutral isosteric analogue 3,3-dimethylbutyraldehyde (DMBA) showed 13,500-fold loss of catalytic efficiency (Table 1). It is noteworthy to mention that DMBA exists mainly as the aldehyde form (≥ 65%) in aqueous solution favoring reductase-catalyzed NADPH-dependent reduction,^37^ whereas glycine betaine aldehyde predominantly exists as the hydrated *gem*-diol form (99%) which must then undergo dehydration to unveil the aldehyde functional group for reduction.

**Table 2.** Inhibition study of ATRR.

| Inhibitor | Enzyme construct | Substrate | Inhibition type | $K_i$ (mM) |
| --- | --- | --- | --- | --- |
| | ATR<sub>1</sub>R<sub>2</sub> | Glycine betaine | Mixed | $K_i=0.27 \pm 0.03$, $K_i'=6.7 \pm 0.9$ |
| Choline | R<sub>2</sub> | Glycine betaine aldehyde | Mixed | $K_i=4 \pm 1$, $K_i'=8 \pm 3$ |
| N(CH<sub>3</sub>)<sub>4</sub><sup>+</sup>Cl<sup>-</sup> | ATR<sub>1</sub>R<sub>2</sub> | Glycine betaine | Competitive | $0.069 \pm 0.007$ |
| | R<sub>2</sub> | Glycine betaine aldehyde | Mixed | $K_i=5 \pm 1$, $K_i'=18 \pm 4$ |
| N(CH<sub>3</sub>)<sub>3</sub>H<sup>+</sup>Cl<sup>-</sup> | ATR<sub>1</sub>R<sub>2</sub> | Glycine betaine | Mixed | $K_i=2.6 \pm 0.5$, $K_i'=53 \pm 4$ |
| N(CH<sub>3</sub>)<sub>2</sub>H<sub>2</sub><sup>+</sup>Cl<sup>-</sup> | ATR<sub>1</sub>R<sub>2</sub> | Glycine betaine | Competitive | $21 \pm 2$ |
| NH<sub>4</sub><sup>+</sup>Cl<sup>-</sup> | ATR<sub>1</sub>R<sub>2</sub> | Glycine betaine | No inhibition | No inhibition up to 1 M inhibitor concentration |

Since the product choline consists of the same head-group, we next tested whether choline could act as an inhibitor competing with glycine betaine and glycine betaine aldehyde. As a result, choline was shown to be a mixed inhibitor for both glycine betaine reductase and aldehyde reductase activity, but more potent (*K*_i_ = 0.27 mM, *K*_i_’ =6.7 mM) in competition against glycine betaine in the first reduction step. Therefore, choline could act as a feedback inhibitor to regulate ATRR enzymatic activity (Figure 3D). The lowered binding affinity of choline to R_2_ (*K*_i_ = 4 mM) favors the release of choline after glycine betaine aldehyde reduction to avoid direct product inhibition.

### Physiological role of ATRR in fungi

Our discovery of ATRR as a glycine betaine reductase strongly indicates that it is the missing link in choline-glycine betaine metabolism (Figure 4). Oxidation of choline to glycine betaine initiates choline catabolic process. This common choline degradation pathway has been found in all domains of life, and is also considered to be irreversible.^38^ Conversion of glycine betaine to choline was proposed as an alternative pathway to choline in fungi, but the underlying biochemical mechanism and genetic basis have remained elusive.^39,40^ A plausible mechanism for this conversion was postulated based on the reduction of L-aspartate to L-homoserine, in which the carboxy group is first activated via phosphorylation by a kinase. This is followed by conversion to a thioester that is reduced into aldehyde by dehydrogenase and then reduced to alcohol by a separate reductase. The biosynthetic logic of ATRR discovered in this work also uses a carboxylic activation strategy for reduction but is more concise as all steps are performed by one enzyme and the key thioester intermediate is channeled through the phosphopantetheine arm of a thiolation domain.

**Figure 4.**
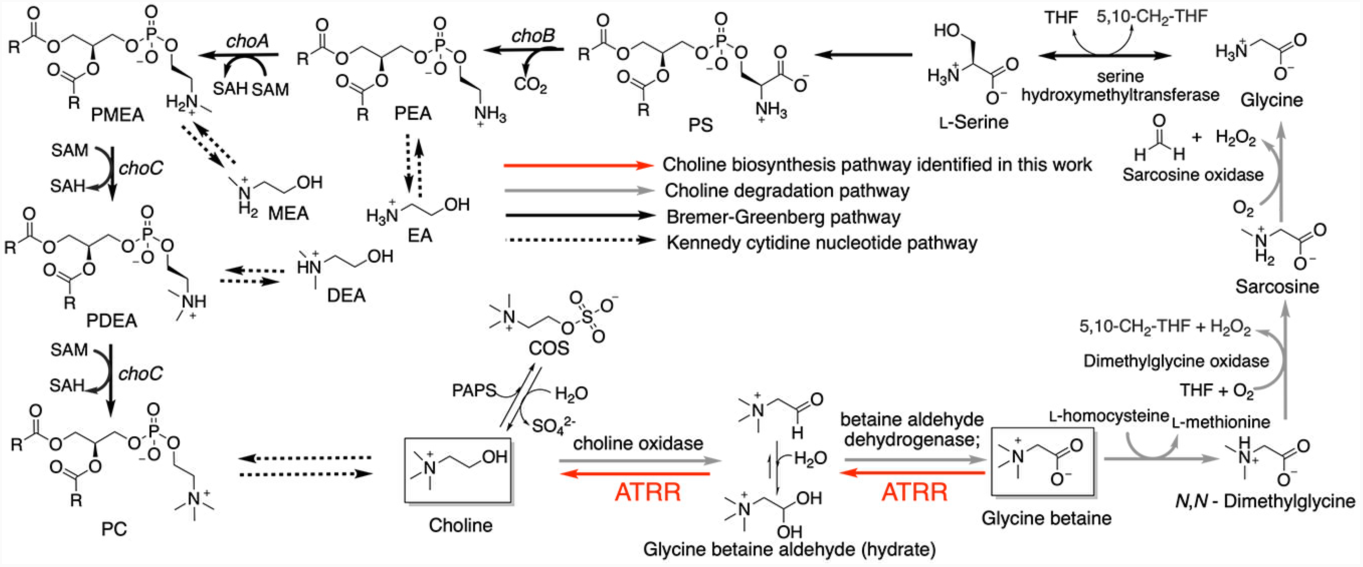
Choline-glycine betaine metabolism in fungi. Abbreviations: PS, phosphatidylserine; PEA, phosphatidylethanolamine; PMEA, phosphatidylmonomethylethanolamine; PDEA, phosphatidyldimethylethanolamine; EA, ethanolamine; MEA, monomethylethanolamine; DEA, dimethylethanolamine; R, fatty acyl chain. PAPS, 3’-phosphoadenosine-5’-phosphosulfate; THF, tetrahydrofolate.

To test whether ATRR plays a role in choline biosynthesis in fungi, we deleted *atrr* gene in both the wild-type (*wt*\*) *A*. *nidulans* and *A. nidulans* Δ*choA* strain by using the split-marker method (*SI Appendix*, Fig. S13-14).^41^ Disruption of *choA* gene encoding the phosphatidylethanolamine (PE) *N*-methyltransferase blocks the Bremer-Greenberg methylation pathway from phosphoethanolamine (PEA) and renders the Kennedy pathway from free choline the only option for biosynthesis of the essential phospholipid phosphatidylcholine (PC, Figure 4).^40^ Deletion of *atrr* gene would abolish conversion of glycine betaine to choline, but both Bremer-Greenberg pathway and Kennedy pathway remain functional. Accordingly, we expect no growth defects of Δ*atrr* strain, while the Δ*choA* strain would show choline/betaine dependent growth. Without the ability to convert glycine betaine to choline, the Δ*atrr*Δ*choA* double knockout strain is expected to be truly choline-auxotrophic even in the presence of glycine betaine. Indeed, when cultured on solid minimal medium (MM), no major growth and morphological difference was observed between Δ*atrr* and *wt\** strain. On the contrary, neither Δ*choA* nor Δ*atrr*Δ*choA* strain was able to grow because of obstructed PC biosynthesis (Figure 5A). The vegetative growth of Δ*choA* strain can be stimulated by supplementation of either choline or glycine betaine (2-20 µM), though full restoration of the conidiation requires higher concentration (>100 µM). In contrast, the Δ*atrr*Δ*choA* strain exhibits choline-auxotroph character: glycine betaine can no longer replace choline to restore growth and asexual development under choline-limiting condition. These results confirm that ATRR rescues the phenotype of Δ*choA* by converting glycine betaine to choline. Therefore, ATRR mediated reduction of glycine betaine is a “shortcut” pathway to choline.

**Figure 5.**
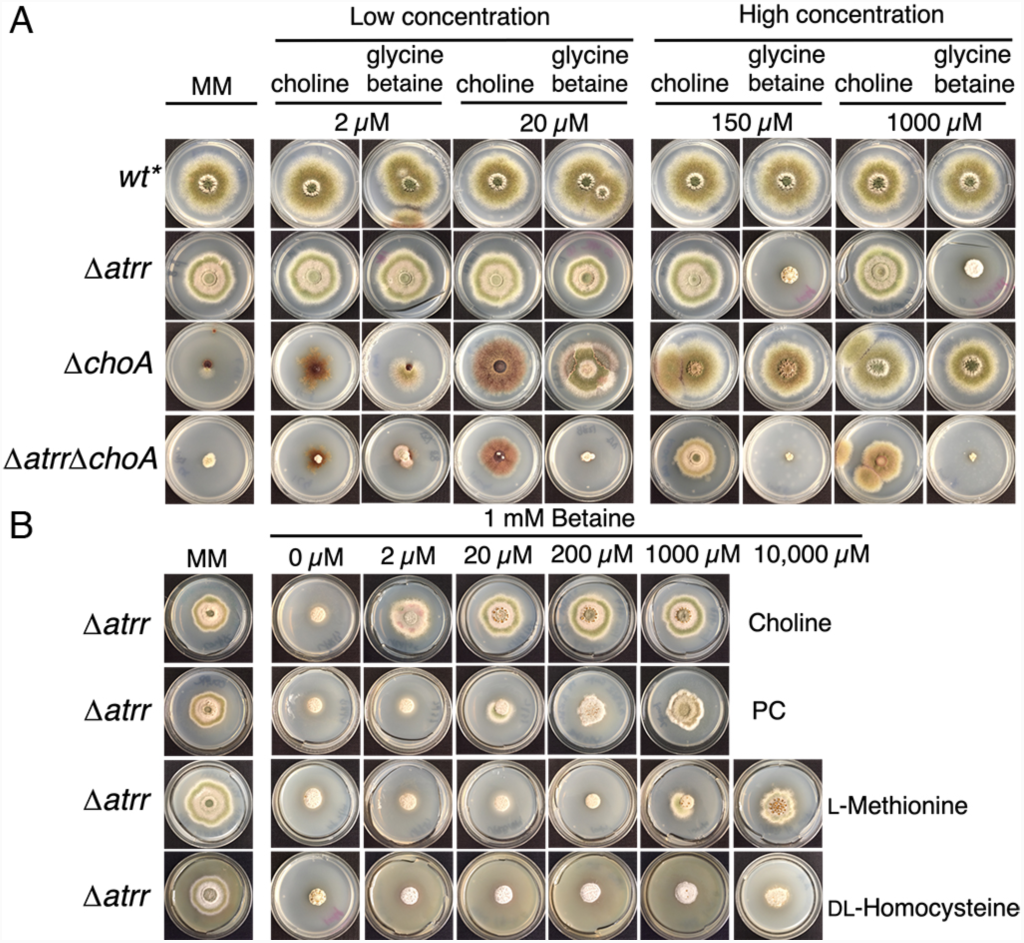
Disruption of *A. nidulans atrr* gene impairs conversion of glycine betaine to choline. *(A*) Growth of *A*. *nidulans* gene deletion mutants on solid minimal medium (MM) at 37 °C for 3 days. The medium was supplemented with different concentrations of glycine betaine and choline. (*B*) Chemical complementation restores growth of *A*. *nidulans* Δ*atrr* strain at high concentration of glycine betaine (1 mM).

Unexpectedly, although no significant difference was observed between the Δ*atrr* and *wt*\* strain growing at low concentration (2-20 µM) of exogenous glycine betaine, the hyphae growth and conidiation of Δ*atrr* strain were severely impaired at high level glycine betaine (>100 µM, Figure 5A), which indicates ATRR function is essential to *A*. *nidulans* under such condition. To interrogate the role of glycine betaine in the absence of ATRR, we co-supplemented choline to the medium. With increasing amount of choline (>20 µM) added, the growth of Δ*atrr* strain can be fully reverted to the normal state even in the presence of millimolar level of glycine betaine (Figure 5B). Similar but not identical results were also observed when PC and L-methionine (10 mM) were co-supplemented. These chemical complementation results suggest the impaired growth caused by high level of exogenous glycine betaine is presumably due to impeded Bremer-Greenberg methylation pathway which again led to deficient PC biosynthesis. High level of methionine could increase SAM biosynthesis via methionine catabolism, which may further up-regulate and drive the methylation pathway against glycine betaine. In comparison, co-supplementation of L-homocysteine (as a racemic L-and D-mixture) in order to favor glycine betaine degradation did not alleviate the impaired growth of Δ*atrr* strain, which suggests the glycine betaine degradation pathway is likely not playing a major role in this phenotype. Nevertheless, inhibition of Bremer-Greenberg methylation pathway by glycine betaine necessitates the function of ATRR in fungi, which not only can supply choline for PC biosynthesis via Kennedy pathway, but also help to ‘detoxify’ high levels of glycine betaine. Therefore, our results confirmed an important physiological role of ATRR in choline and PC metabolism in *A. nidulans*; and also reveal a previously unknown effect of glycine betaine to *A. nidulans* growth.

## DISCUSSION

Our study here was driven by a highly conserved NRPS-like enzyme in fungi with an unusual domain architecture. Successful elucidation of its unknown function as glycine betaine reductase serendipitously led to the resolution of a long-standing question in choline-glycine betaine metabolism, whether the common choline-glycine betaine pathway is reversible. It is now clear that this pathway can run in the reverse direction, and our study provides the molecular basis: ATRR catalyzes ATP-and NAPDH-dependent reduction of glycine betaine by a catalytic mechanism similar to that of well-known NRPS-dependent CARs, generating glycine betaine aldehyde intermediate, which is further reduced at the R_2_ domain unique to ATRR, yielding final product choline. Orthologues of ATRR are found to be prevalent in fungi and exhibit high sequence identities with characterized *A. nidulans* ATRR (*SI Appendix*, Fig. S2-3), which suggests that this efficient, single-enzyme catalyzed reductive choline biosynthesis may be widespread in fungi. Harboring this enzyme permits direct reutilization of endogenously stored glycine betaine (∼3 mM) for on-demand biosynthesis of choline and choline-derivatives, including phospholipid PC (essential role in maintaining membrane integrity and functionality), and choline-*O*-sulfate (a mean for intracellular sulfate storage).^42^ Comparing to the high metabolic cost of the *de novo* choline biosynthetic pathway (Bremer-Greenberg pathway) at the expense of three consecutive methylation steps (costing 12 ATP molecules per methylation event),^43^ regeneration of choline from glycine betaine is much more cost-effective (costing 1 ATP and 2 NADPH). More importantly, this “shortcut” pathway also acts as an “emergency safeguard” pathway to supply choline from its “reserve form” glycine betaine for PC biosynthesis when the Bremer-Greenberg methylation pathway is obstructed.

Uncovering ATRR function was rooted in understanding the biochemical logic of NRPS-like CARs. We reasoned that A domain must play a gatekeeping role in selecting and activating carboxylic acid substrate; and identifying the substrate would reveal the function of ATRR. Although our initial search for the substrate guided by the 10 AA code was unsuccessful, key insights of A domain substrate specificity was provided by our interpretation of the structure-function relationship based on a simple homology model, which prioritized a pool of potential substrates and the long-sought after substrate glycine betaine was identified eventually by experimental enzymatic activity screening. Our results expanded the biochemistry scope of NRPS family: ATRR-A is shown to be the first adenylation domain that can activate and thioesterify a betaine-type substrate; and convincing evidence supports our hypothesis that cation-π play an important role in substrate recognition at the adenylation domain, which for the first time implicates cation-π interaction in the NRPS family enzymology. Our findings support the notion that cation-π interaction is a fundamental and universal non-covalent interaction for substrate/ligand recognition in biology.^26,27^

In summary, our study demonstrated that the structure-guided approach can be a general strategy for functional discovery of uncharacterized NRPS-like enzymes, and orphan NRPS assembly lines from the ever-increasing size of genomic data, which may aid discovery of new metabolic pathways and natural products.^11^ In principle, such process can be greatly facilitated by docking a library of carboxylic acid metabolites to high quality homology models of adenylation domains.

## MATERIALS AND METHODS

Materials and methods are summarized in SI Appendix, including Figs. S1–S14 and Tables S1.

## ACKNOWLEDGMENT

This work was supported by the NIH (1R35GM118056) to Y.T.. Y.H. is a Life Sciences Research Foundation fellow sponsored by the Mark Foundation for Cancer Research. We thank Dr. Mengbin Chen for helpful discussions, and Abigail Yohannes for helping with bioinformatic analysis.

