## Supplementary Information for "Structure-guided function discovery of an NRPS-like glycine betaine reductase for choline biosynthesis in fungi"

This PDF file includes:

Supplementary Table

Supplementary Figures

### 1. Materials and Methods

#### 1.1. Bioinformatics

We used *A. nidulans atrr* (AN5318.2) protein sequence as a query to retrieve nonredundant *atrr* orthologues by using blastp and tblastn. As a result, a complete dataset of 322 *atrr* orthologues were obtained. The criteria we used to filter false positive and redundant hits were: 1) All putative *atrr* orthologues must share the same domain architecture (A-T-$R_1$-$R_2$). 2) only one *atrr* orthologue is collected per species. In most species, only one *atrr* gene is present, except that in *Trichoderma citrinoviride*, *Hortaea werneckii* EXF-6669, and *Rachicladosporium* sp. CCFEE 5018 two nearly identical *atrr* paralogues (>96% amino acid sequence identity) are found in each genome.

For genome annotation, 2ndFind program was used to predict the open reading frame and intron, gene function was assigned based on BlastP search. To annotate the genomic context, contigs containing *atrr* gene were found by tblastn and opened using Snapgene Viewer version 1.5.3. Onine programs, such as 2ndFind, Softberry, and blastx, were used to find surrounding genes. Function of gene products was assigned by blastp of each protein's predicted amino acid sequence and examination of putative conserved domains.

To analyze the nonribosomal code of ATRR-A, we performed multiple sequence alignment of selected adenylation domain, the 10AA code of each sequence was manually identified based on the study of PheA.[1,2] The 10-aa code of ArbA was directly from ref. 3. The accession number for the rest sequences analyzed in Figure 2a are shown in paratheses here: GrsA-PheA (P0C061), AcvA (P27742),[4] SidN (K7NCP5),[5] ClbH (Q0P7J8),[6] AdmJ (Q84FL3),[7] SgcC1 (AAL06681),[8] BecJ (FJ872523.1),[9] Lys2 (P07702),[10] ATRR (AN5318.2), AnaPS (A1DN09),[11] TomA (ACN39014),[12] DhbE (AAN15214.1),[13] MbtA (YP_0093597501.1),[14] ATEG_03630,[15] TdiA (ABU51602.1),[16] RalA (AEC03968.1),[17] AtrA (AUO29225.1).[18] Note that the following sequences were not included in the A domain analysis due to poor alignment with other ATRRs, indicating different functions: *Sphaeroforma arctica* JP610 (XP_014150094.1), *Wuchereria bancrofti* (EJW86348.1), *Lingul anatine* (XP_013419834.1)

#### 1.2. Chemicals and general methods

Glycine betaine aldehyde and stachydrine are purchased from Cayman Chemicals. and. β-Alanine betaine was purchased from Chemspace. Butyrobetaine, L-hercynine, ergothionine, and sulfobetaine were purchased from Toronto Research Chemicals. L-α-phosphatidylcholine (95%) (Soy) was purchased from Avanti Polar Lipids, Inc. Isopropyl-β-D-1-thio-galactopyranoside (IPTG) was purchased from Carbosynth. Tris-(2-carboxyethyl) phosphine hydrochloride (TCEP-HCl) was purchased from GoldBio Biotechnology. All other chemicals were purchased from Sigma-Aldrich. PCR reactions were performed using the Phusion® high-fidelity DNA polymerase (New England Biolabs) and used according to the manufacturer's instructions. Custom oligonucleotides were synthesized by Integrated DNA Technologies. *Escherichia coli* strain DH10B was used for cloning procedures.

#### 1.3. Protein expression and purification

The *atrr* gene (AN8150.2) can be cloned from the genomic DNA extract of *A. nidulans* ΔEM strain.[19] Different expression constructs were made by subcloning the corresponding domain region into a modified pET28a (+) vector (Addgene plasmid #29656). The resulting N-terminal hexa-histidine tagged proteins were overexpressed in *E. coli* BL21(DE3) in LB medium in the presence of 50 mg/L kanamycin. Expression was induced by 100 μM IPTG when $OD_{600}$ reached 1.0, and cell cultures were left grown at 16 °C overnight. Cells were harvested by centrifugation and resuspended in cell lysis buffer [50 mM $K_2HPO_4$ (pH 8.0), 300 mM NaCl, 10% (v/v) glycerol]. Cells were lysed by sonication and the cell lysate was cleared by centrifugation at 26,000 g for 30 min at 4 °C. The supernatant was incubated with $Ni^{2+}$-NTA resin for 2 hrs and then the slurry was loaded onto a gravity column. The resin was washed and eluted with increasing concentrations of imidazole in cell lysis buffer. The fractions were examined by SDS-PAGE gels and targeted proteins were subject to size-exclusion chromatography by using a HiLoad Superdex 200 26/60 column (GE Healthcare) equilibrated in storage buffer [50 mM HEPES (pH 7.5), 150 mM NaCl, 1 mM TCEP, 10% (v/v) glycerol]. Pure fractions were concentrated to 10 mg/mL by Amicon concentrators (Millipore) and stored at -80 °C. Protein concentrations were determined by Bradford assay.

The majority of ATRR proteins purified from *E. coli* are in the apo-form. To enzymatically convert it to its functional holo-form, before size-exclusion chromatography, apo ATRR proteins (at concentration ~10 mg/mL) were incubated with in-house purified phosphopantetheinyl transferase NpgA (0.1 mg/mL final concentration), 10 mM $MgCl_2$, 500 μM CoA on ice for 2 hours.[20]

ATRR mutants and domain truncation variants are purified similarly to the wild-type.

#### 1.4. Substrate screening in vitro

The hydroxylamine-trapping assay detecting adenylation activity was performed according to the protocol described in ref. 21. Briefly, the reaction was initiated by mixing 150 μL of substrate mixture [50 mM Tris, (pH 8.0), 30 mM $MgCl_2$, 300 mM hydroxylamine (pH 8.0), 10 mM carboxylic acid substrate] with equal volume of enzyme mixture [100 mM Tris (pH 8.0), 20 mM ATP, 20 μM enzyme (full-length ATRR or excised A domain)]. For some hydrophobic substrates, 2-5% (v/v) DMSO was included to facilitate dissolving the substrate. The reaction mixture was then incubated at 30 °C for 16 hrs. The reaction was stopped by mixing with 300 μL of stopping solution [10% (w/v) $FeCl_3$•$6H_2O$ and

3.3% TCA dissolved in 0.7 M HCl]. The precipitated enzymes were removed by centrifugation at 17,000 g for 5 min, and 200 μL of the supernatant were transferred to a 96-well plate and the absorbance of the ferric-hydroxamate complex at 540 nm was measured by using a Tecan M200 plate reader.

The carboxylic acid reductase activity was performed according to the protocol described in ref. 22. Briefly, 50 μL of reaction mixture [100 mM HEPES, pH 7.5, 10 mM ATP, 15 mM $MgCl_2$, 1 mM NADPH, 10 mM carboxylic acid substrate] was mixed with equal volume of enzyme mixture [100 mM HEPES, pH 7.5, 20 μM ATRR] to initiate the reaction. The reductase activity was monitored by following the oxidation of NADPH at 340 nm.

#### 1.5. Steady-state kinetics measurement

For the steady-state kinetics measurement, initial rates were taken from the progress curve of the continuous assay based on NADPH consumption. A standard curve of NADPH absorbance at 340 nm was constructed. The steady-state kinetics of aldehyde reductase was performed similarly, except without addition of ATP and $MgCl_2$. The kinetic parameters were obtained by fitting the data to the following equations:

$$v = \frac{V_{max}\,[S]}{K_m + [S]}$$

Substrate inhibition:

$$v = \frac{V_{max}\,[S]}{K_m + [S](1 + \frac{[S]}{K_i})}$$

For inhibition study, an IC50 assay was first performed to get preliminary data of the inhibitor dissociation constant $K_i$ in order to define the range varying inhibitor concentration. Steady-state kinetics experiments were then performed in the presence of different concentration of inhibitors. A Line-weaver burke plot was constructed to determine the inhibition type. The inhibitor dissociation constant $K_i$ was obtained by fitting the steady-state kinetics globally with the following equation:

Competitive inhibition:

$$v = \frac{V_{max}\,[S]}{K_m(1 + \frac{[I]}{K_i}) + [S]}$$

Mixed inhibition:

$$v = \frac{V_{max}\,[S]}{K_m(1+\frac{[I]}{K_i}) + [S](1+\frac{[I]}{K_i'})}$$

### 1.6. Biochemical characterization of intermediate during glycine betaine reduction.

The glycine betainoyl thioester intermediate was characterized by using mass-spectrometer. Purified stand-alone T domain was first converted to holo-form. Holo-T domain was then loaded with glycine betaine by co-incubation with A domain. Briefly, 200 µL of reaction mixture [50 µM of T domain, 500 µM A domain, 10 mM $MgCl_2$, 10 mM ATP, 20 mM glycine betaine] was incubated at 30 °C for 30 min. The reaction mixture was desalted by using the Zeba spin desalting column (Thermo Fisher Scientific). The acyl-intermediate of T domain was immediately characterized by MALDI-TOF (Bruker Ultraflex). Before analysis, the sample was serial diluted with water, and mixed with equal volume of sinapic acid matrix. Prolonged incubation of the reaction mixture (1-2 hrs) did not increase the occupancy of gycine betainoyl moiety, presumably due to the instability of the thioester since longer incubation (>2 hrs) led to less occupancy.

The aldehyde intermediate was characterized by continuous trapping with phenylhydrazine. Basically, 10 mM freshly made phenylhydrazine was inclused in the reaction mixture. The phenylhydrazone derivative was separated by LC-MS, performed on a Shimadzu 2020 EVLC–MS (Phenomenex kinetex, 1.7 µm, 2.0 x 100 mm, C18 column) using positive and negative mode electrospray ionization with a linear gradient of 5−95% MeCN−H2O supplemented with 0.1% (v/v) formic acid in 15 min followed by 95% MeCN for 5 min with a flow rate of 0.3 mL/min. The identity of the intermediate was confirmed by comparing the retention time with derivatized standard.

The final product choline accumulated in the reaction mixture was characterized by derivatization with 1-naphthyl isocyanate as described in ref. 23, and the derivatives were analyzed by using LC-MS.

### 1.7. Genetic manipulation

The deletion mutant of *A. nidulans* Δ*atrr* was constructed using a split-marker approach.[24] Briefly, ~1-kb fragments flanking the targeted deletion region of *atrr* were amplified by PCR from *A. nidulans* ΔEM genomic DNA extract, and the *Aspergillus fumigatus pyrG* marker gene was amplified from the plasmid pYTU.[19] The three fragments were joined to create a deletion cassette by using homologous recombination with vector pXW55 in *S. cerevisiae.* The deletion cassette was then spitted into half by PCR amplification with an overlapping region of 500 bp spanning *pyrG* marker. Genes were deleted in parental *A.nidulans* ΔEM strain. Single colonies were picked 3 days after the transformation to avoid cross-contamination. Genomic DNA of picked transformants were isolated and gene deletion was confirmed by PCR and Sanger DNA sequencing of PCR fragment (Figure S12).

The *A. nidulans* Δ*choA* mutant was constructed similarly, except that *riboB* marker was used and the gene fragment was PCR amplified from plasmid pYTR. The double deletion mutant *A. nidulans* Δ*atrr*Δ*choA* was constructed sequentially based on mutant strain *A. nidulans* Δ*atrr*. Gene deletion was confirmed by PCR and Sanger DNA sequencing of PCR fragment (Figure S12).

Strains were maintained as glycerol stocks and activated on solid minimal medium (MM, 10 g glucose, 6 g $NaNO_3$, 0.52 g KCl, 0.52 g $MgSO_4 \cdot 7H_2O$, 1.52 g $KH_2PO_4$, 1 x trace elements, 20 gram agar) at 37 $^oC$ with appropriate supplements (uracil, pyridoxine, or riboflavin). For choline-auxotroph mutant strains Δ*choA* and Δ*atrr*Δ*choA*, 1 mM choline was supplemented in the medium.

### 2. Figures

#### 2.1. Cladistic summary of *atrr* genes distribution

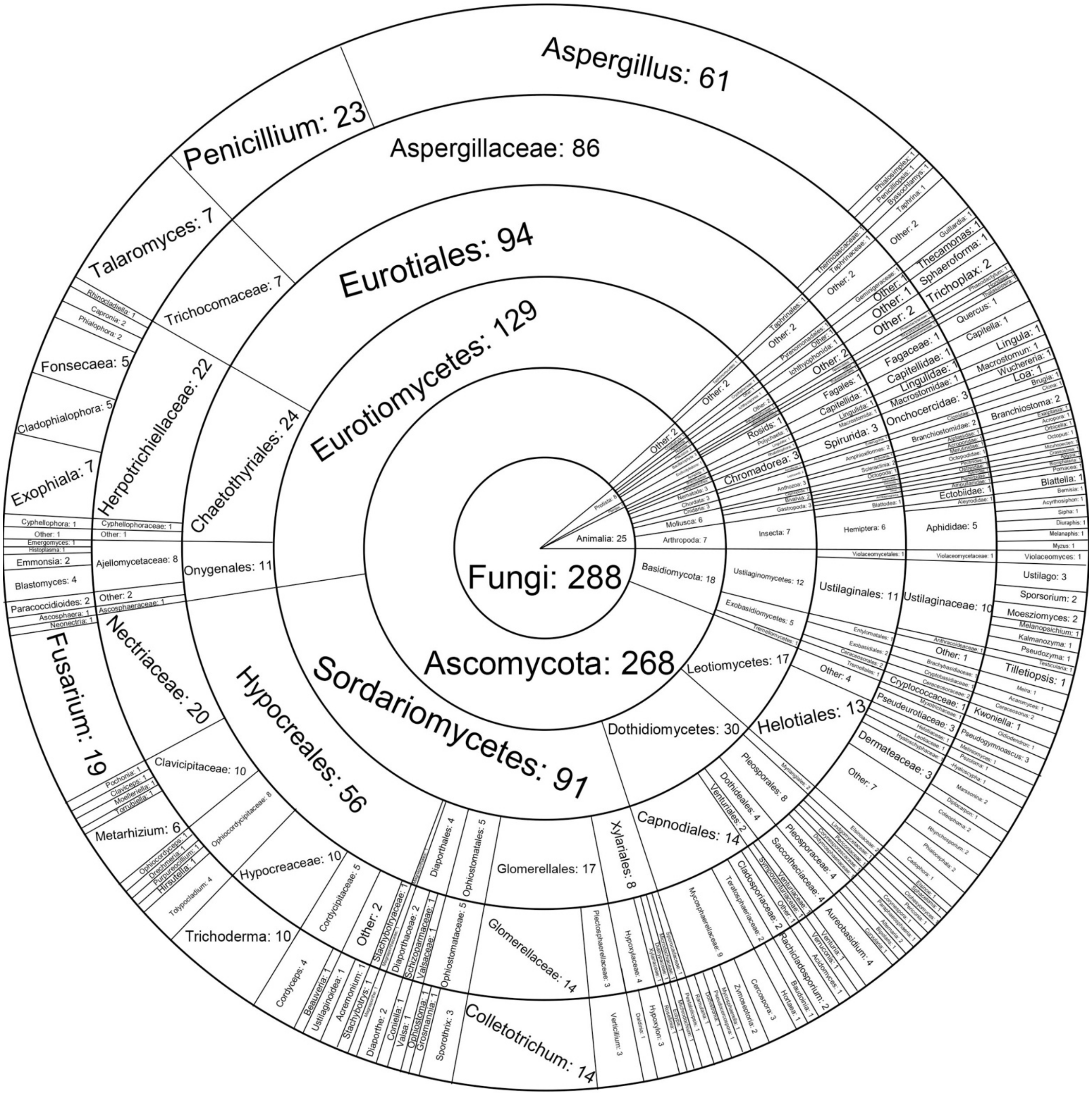

Fig. S1. Cladistic summary of all ATRR orthologues in sunburst format (from kingdom to genus). Each orthologue's cladistic information was found from its NCBI protein page, and organization of cladistic categories was done with reference to NCBI taxonomy. Orthologues originating from an organism which is *incertae sedis* within a taxonomic rank are grouped into "other", as are those which do not have one or more of the six taxonomic levels. The vast majority of orthologues exist in *Fungi* (288), with a small number in *Animalia* (25), *Protista* (8), and *Plantae* (1).

### 2.2. Genomic context analysis of *atrr* orthologues from different fungal genera.

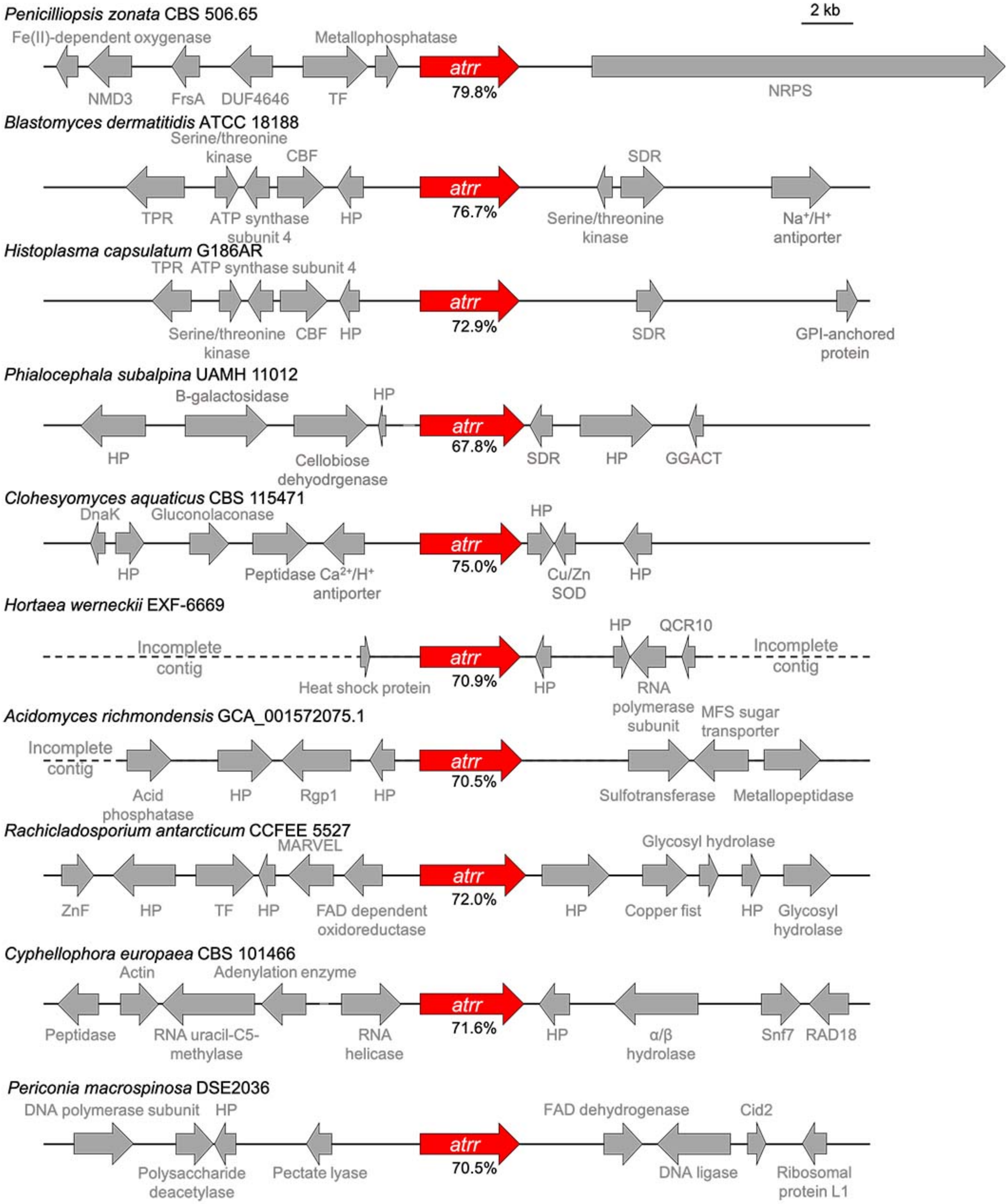

Fig. S2. Continued on next page.

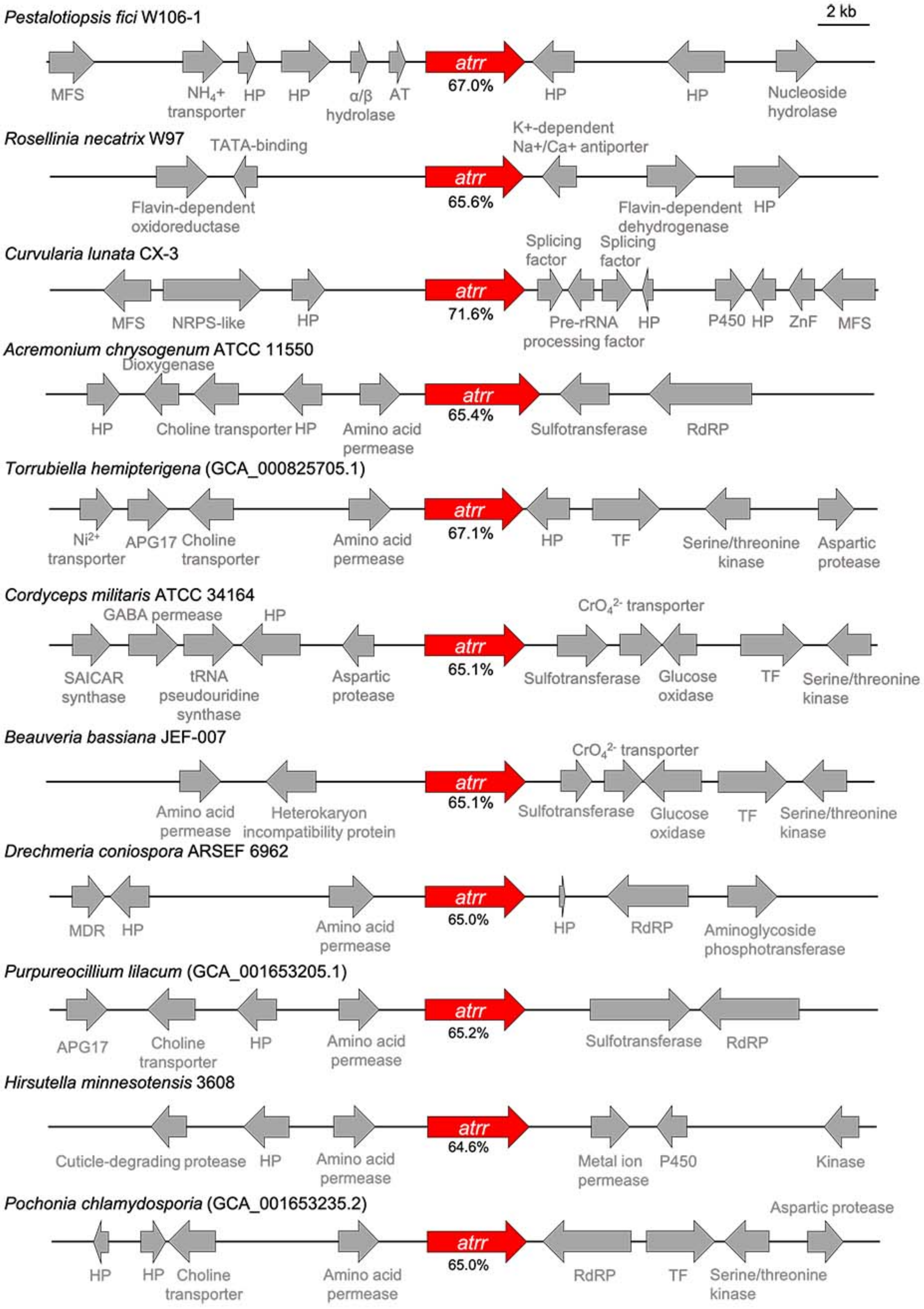
Pestalotiopsis fici W106-1
2 kb
atrr
67.0%
MFS
NH4+ transporter
HP
HP
α/β hydrolase
AT
HP
HP
Nucleoside hydrolase
Rosellinia necatrix W97
TATA-binding
K+-dependent Na+/Ca+ antiporter
atrr
65.6%
Flavin-dependent oxidoreductase
Flavin-dependent dehydrogenase
HP
Curvularia lunata CX-3
Splicing factor
Splicing factor
atrr
71.6%
MFS
NRPS-like
HP
Pre-rRNA processing factor
HP
P450
HP
ZnF
MFS
Acremonium chrysogenum ATCC 11550
Dioxygenase
atrr
65.4%
HP
Choline transporter
HP
Amino acid permease
Sulfotransferase
RdRP
Torrubiella hemipterigena (GCA_000825705.1)
atrr
67.1%
Ni2+ transporter
APG17
Choline transporter
Amino acid permease
HP
TF
Serine/threonine kinase
Aspartic protease
Cordyceps militaris ATCC 34164
GABA permease
HP
CrO42- transporter
atrr
65.1%
SAICAR synthase
tRNA pseudouridine synthase
Aspartic protease
Sulfotransferase
Glucose oxidase
TF
Serine/threonine kinase
Beauveria bassiana JEF-007
CrO42- transporter
atrr
65.1%
Amino acid permease
Heterokaryon incompatibility protein
Sulfotransferase
Glucose oxidase
TF
Serine/threonine kinase
Drechmeria coniospora ARSEF 6962
atrr
65.0%
MDR
HP
Amino acid permease
HP
RdRP
Aminoglycoside phosphotransferase
Purpureocillium lilacum (GCA_001653205.1)
atrr
65.2%
APG17
Choline transporter
HP
Amino acid permease
Sulfotransferase
RdRP
Hirsutella minnesotensis 3608
atrr
64.6%
Cuticle-degrading protease
HP
Amino acid permease
Metal ion permease
P450
Kinase
Pochonia chlamydosporia (GCA_001653235.2)
Aspartic protease
atrr
65.0%
HP
HP
Choline transporter
Amino acid permease
RdRP
TF
Serine/threonine kinase

Fig. S2. Continued on next page.

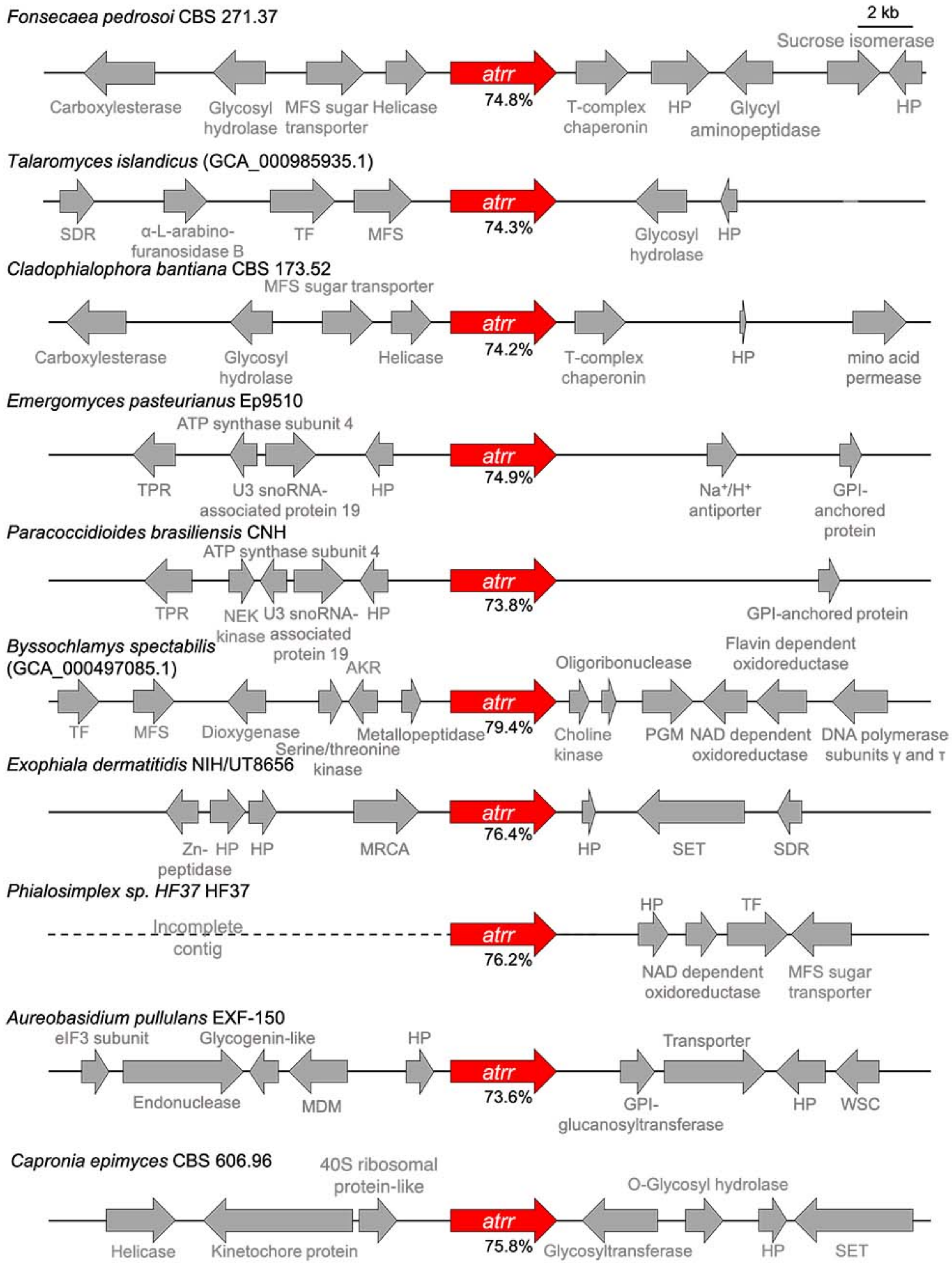

Fig. S2. Continued on next page.

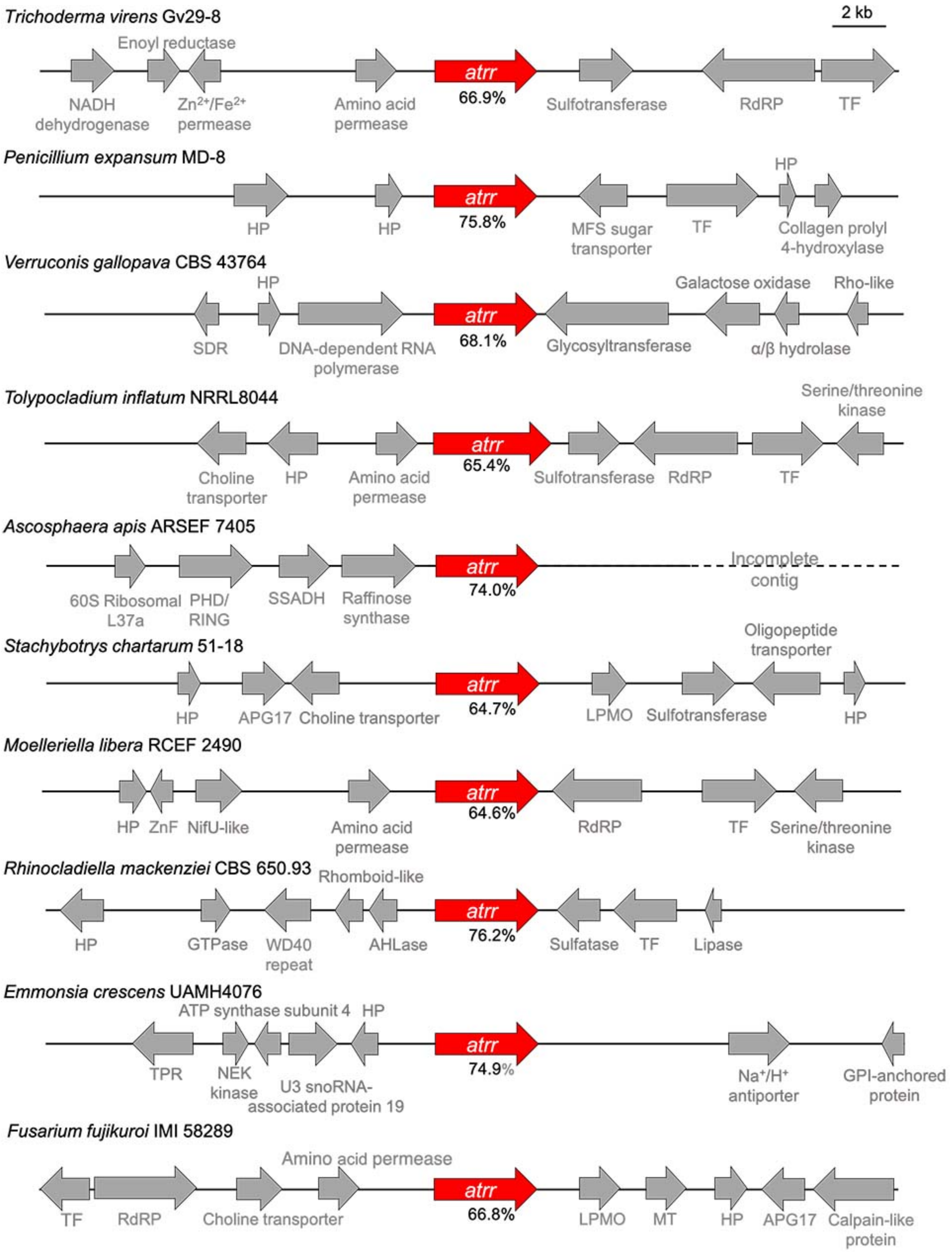

Fig. S2. Continued on next page.

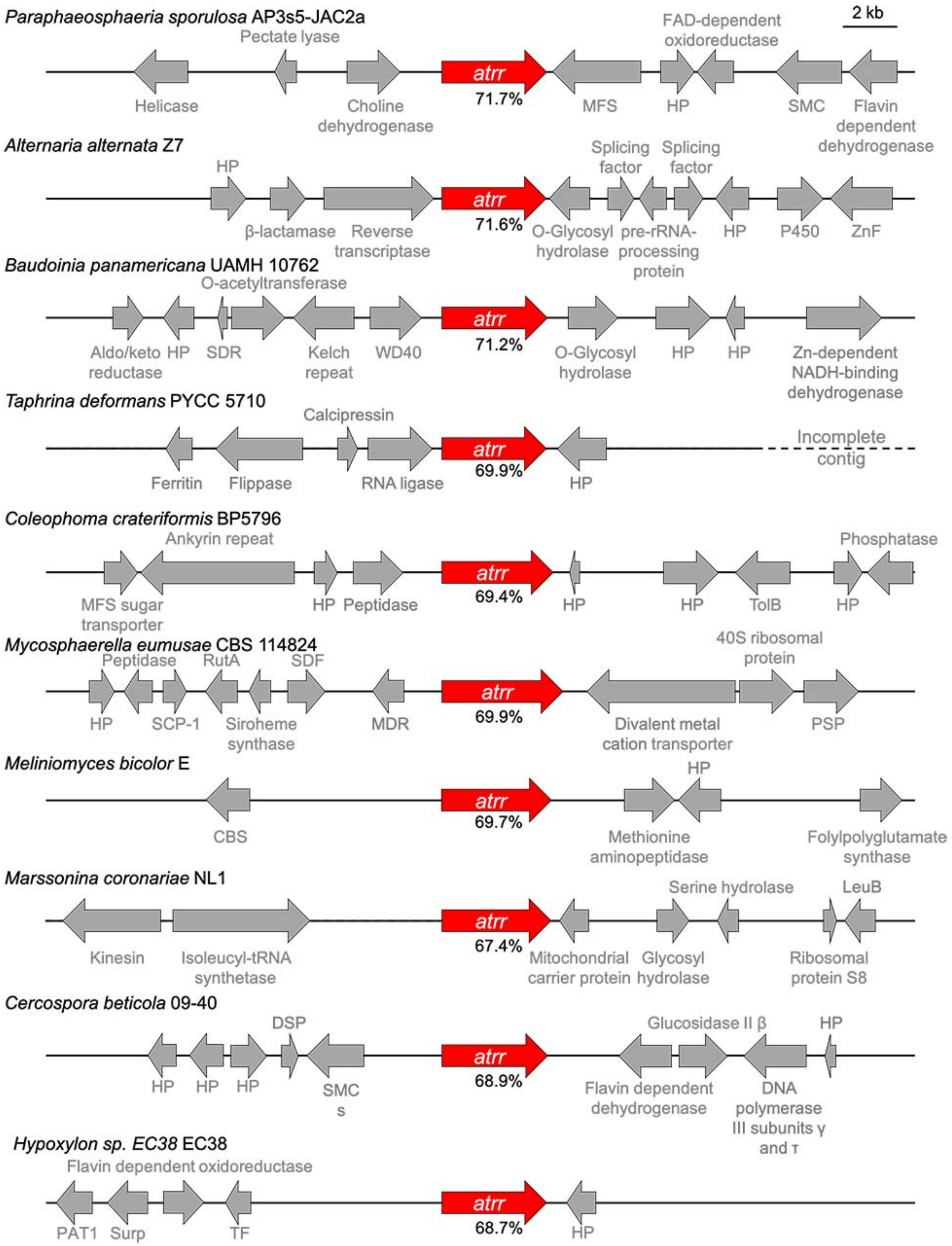

Fig. S2. Continued on next page.

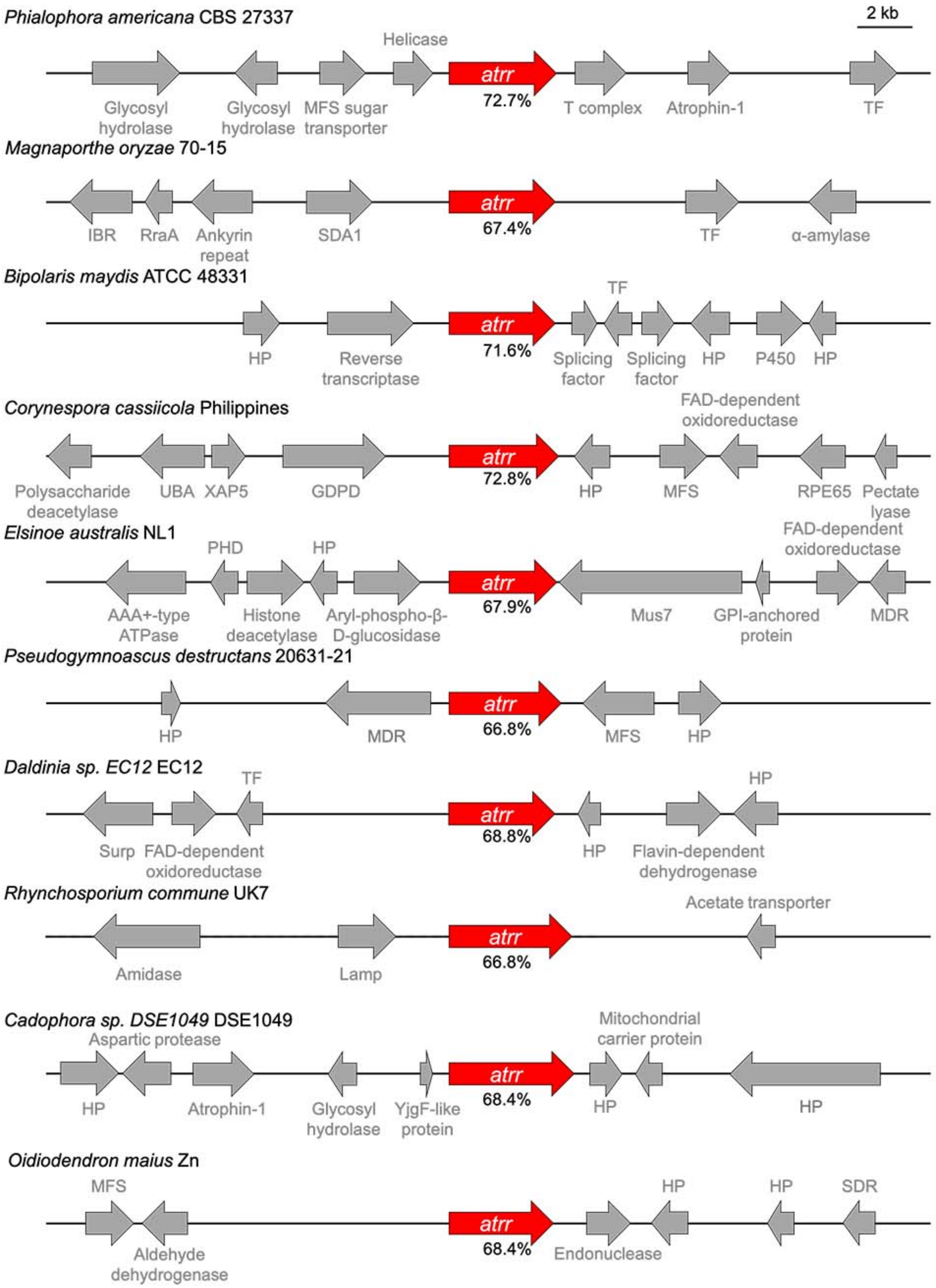

Fig. S2. Continued on next page.

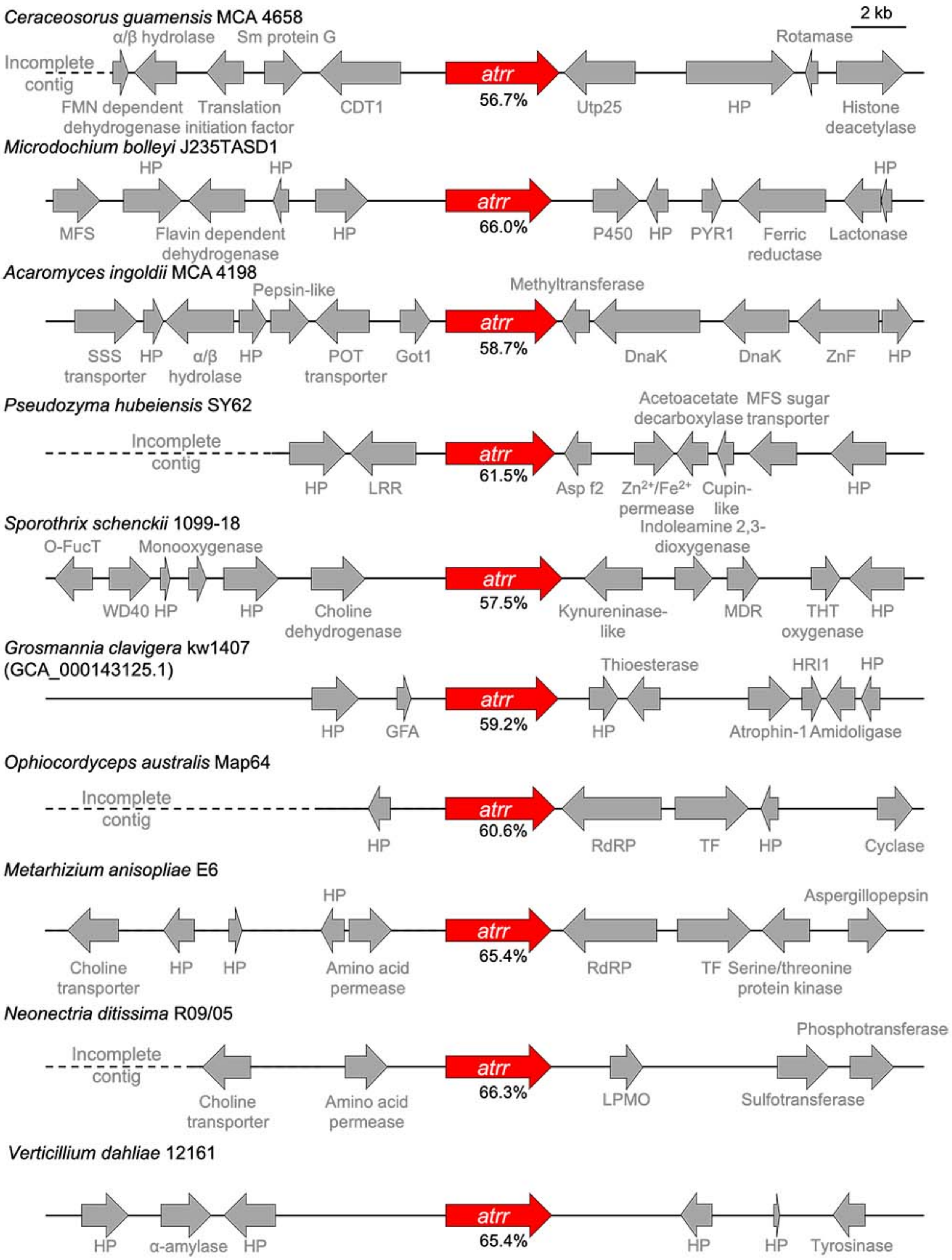

Fig. S2. Continued on next page.

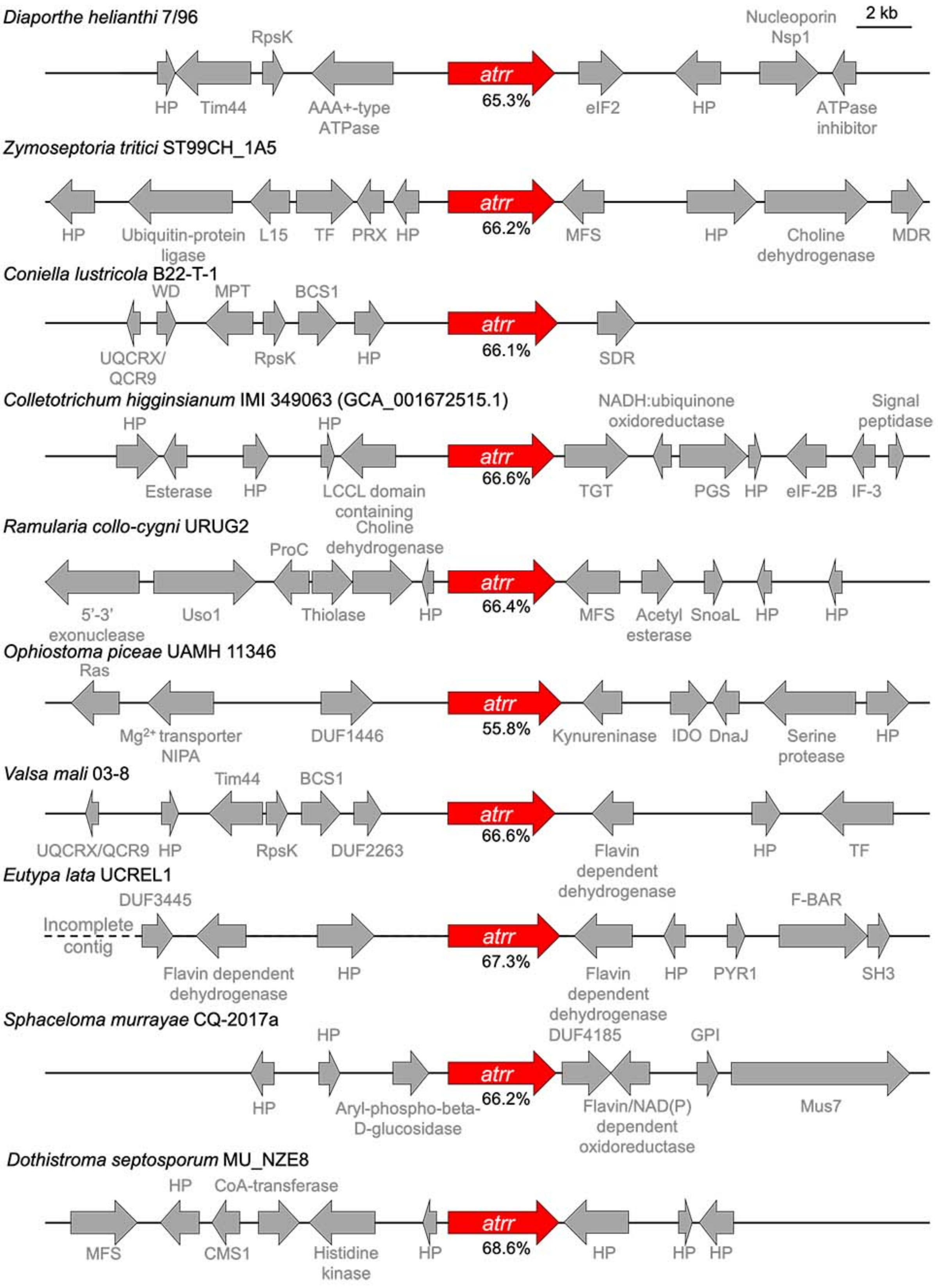

Fig. S2. Continued on next page.

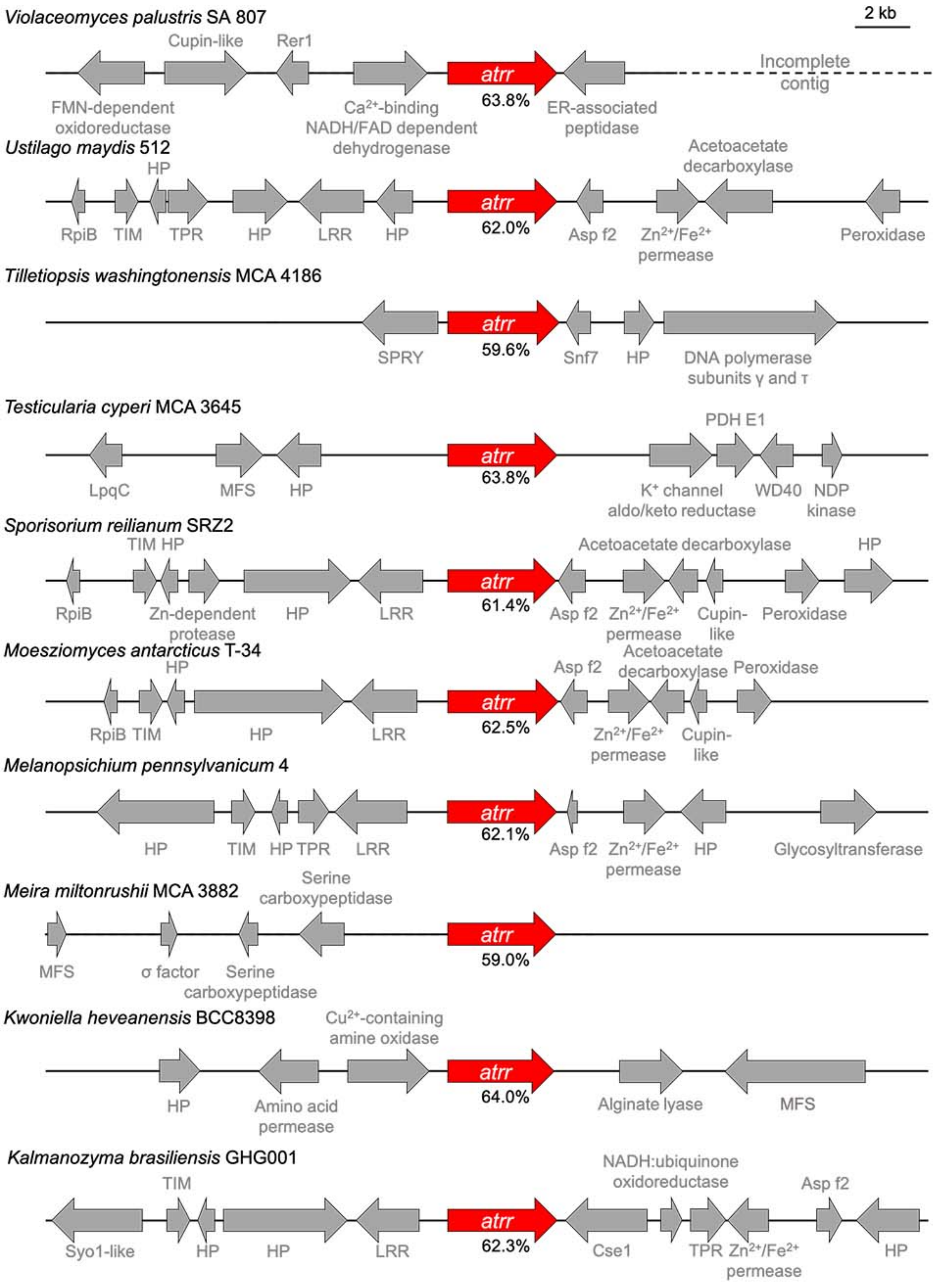

Fig. S2. Continued on next page.

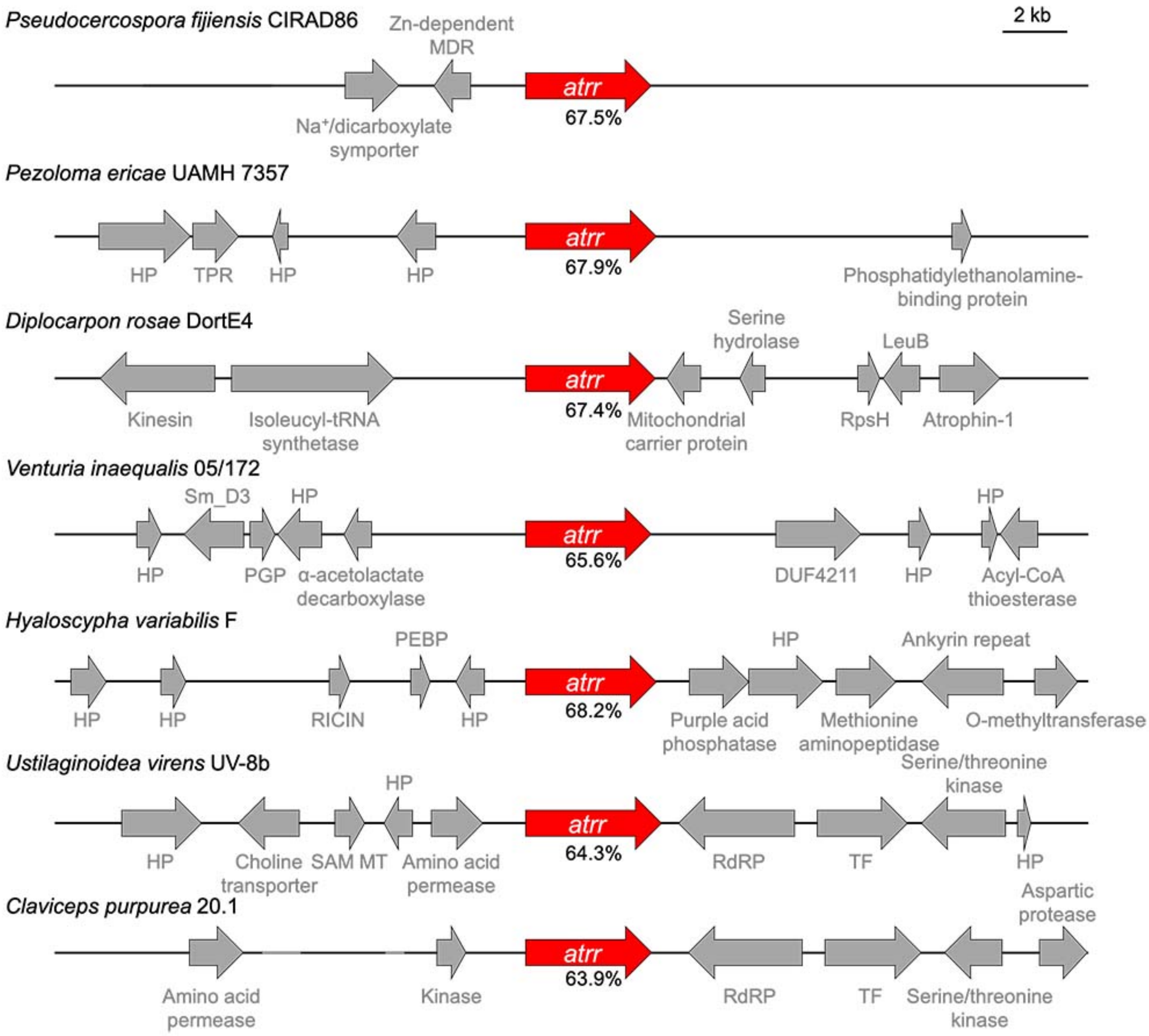

Figure 2. Genomic context analysis of atrr orthologues from representative species for different genera. Numbers under gene arrow indicate the amino acid sequence identity of atrr orthologues relative to A. nidulans atrr. Difference in atrr gene label size is due to introns. *Acremonium chrysogenum* ATCC 11550 had a remarkable 10% intron content. Grey portions of the DNA line indicate Ns in the contig. Abbreviations to follow: HP, hypothetical protein; TF, transcription factor; RdRP, RNA-dependent RNA polymerase (eukaryotic); AT, acetyltransferase; APG17, autophage protein Apg17; SSADH, succinate semialdehyde dehydrogenase; LPMO, lytic polysaccharide monooxygenase; AHLase, N-acyl-homoserine lactonase; WD40, WD40 repeat containing protein; PGM, phosphoglucomutase; AKR, aldoketoreductase; MFS, MFS transporter superfamily; SET, lysine methyltransferase; MRCA, myosin-cross-reactive antigen; MDM, Yeast mitochondrial distribution and morphology; EIF3, translation initiation factor 3 (eIF3) subunit; T complex, TCP-1 chaperonin family; FrsA, fermentation-respiration

switch protein; NMD3, NMD3 family protein; CBF/MAK21, CBF/MAK21 family protein; TPR, tetratricopeptide repeat containing; P450, cytochrome P450; ZnF, zinc finger containing; GGACT, gamma-glutamylamine cyclotransferase; SOD, superoxide dismutase; QCR10, QCR10 family protein; Rgp1, Rgp1 protein; MARVEL, MARVEL domain containing protein; Snf7, Snf7 family protein; rad18, rad18 family protein; Cid2, caffeine-induced death family protein; SMC, structural maintenance of chromosomes family protein; TolB, propeller repeat family protein; PSP, phosphoserine phosphatase; MDR, medium chain reductase/dehydrogenase; SDF, Sodium:dicarboxylate symporter family protein; RutA, RutA-like protein; SCP-1, synaptonemal complex protein 1-like protein; CBS, CBS domain containing; LeuB, isocitrate/isopropylmalate dehydrogenase; DSP, dual specificity phosphatases; Surp, surp module containing (also known as SWAP domain containing) protein; PAT1, topoisomerase II-associated protein family; SDA1, SDA1 homolog; RraA, ribonuclease activity regulator RraA family protein; IBR, IBR domain containing; GDPD, Glycerophosphodiester phosphodiesterase of YPL110cp domain containing protein; XAP5, circardian clock regulator; UBA, UBA domain containing; Mus7, homology to Mus7; PHD, PHD zinc finger containing; AAA+-type ATPase, member of ATPase family associated with various cellular activities (AAA); LAMP, homology to lysosome-associated membrane glycoprotein; YjgF, member of YjgF family; ER-associated peptidase, endoplasmic reticulum associated peptidase; Rer1, Rer1 family protein; Cupin-like, Cupin-like domain; Asp f2, peptidase domain from allergen Asp f2; TIM, triosephosphate isomerase-like; RpiB, ribose 5-phosphate isomerase; SPRY, SPRY domain from Ash2 containing; PDH E1, pyruvate dehydrogenase E1 subunit-like; NDP kinase, nucleoside diphosphate kinase Group I-like; LpqC, poly(3-hydroxybutyrate) depolymerase; Cse1, Cse1 nuclear export receptor domain containing; Utp25, Utp25 family protein; CDT1, CDT1-like; Sm protein G, Sm family protein; Got1, Got1-like; POT transporter, proton-dependent oligopeptide transporter; SSS transporter, solute:sodium symporter family protein; LRR, leucine-rich repeat containing; THT oxygenase, 2,4,5 trihydroxytoluene (THT) oxygenase; O-FucT; O-fucosyltransferase-like; HRI1, HRI1-like; GFA, glutathione-dependent formaldehyde-activating enzyme; PYR1, pyrabactin resistance 1 family protein; eIF2, translation initiation factor 2 subunit gamma-like; Nucleosporin Nsp1, homology to Nsp1; RpsK, homology to ribosomal protein S11; Tim44, homology to Tim44; PRX, peroxiredoxin-like; L15, homology to 60s ribosomal protein L15; BCS1, ATPase BSC1 N terminal domain containing; MPT, mitochondrial protein translocase family protein; WD, uncharacterized protein with Trp-Asp repeat; UQCRX/QCR9, ubiquinol-cytochrome C reductase-like; TGT, queuine tRNA-ribosyltransferase family protein; PGS, homology to CDP-diacylglycerol--glycerol-3-phosphate 3-phosphatidyltransferase; eIF2B: homology to translation inhibitor factor 2B subunit; IF-3, homology to translation initiation factor IF-3; SnoaL, SnoaL fold containing protein; ProC, pyrroline-5-carboxylate reductase; Uso1, homology to Uso1/p115; IDO, indoleamine 2,3-dioxygenase; DnaJ, DnaJ domain containing; $Mg^{2+}$ transporter NIPA, magnesium transporter NIPA-like; Ras, rat sarcoma GTPase family protein; GPI, glycosyl-phosphatidylinositol-anchored protein family; CMS1, U3-containing 90S pre-ribosomal complex subunit; RupsH, ribosomal protein S8-like; PGP, mitochondrial PGP phosphatase; Sm_D3, Sm protein D3; PEBP, phosphatidylethanolamine-binding protein; RICIN, ricin-type beta-trefoil lectin domain containing protein; DUF, domain of unknown function;

#### 2.3. Genomic context analysis of *atrr* orthologues from species of genus *Aspergillus*

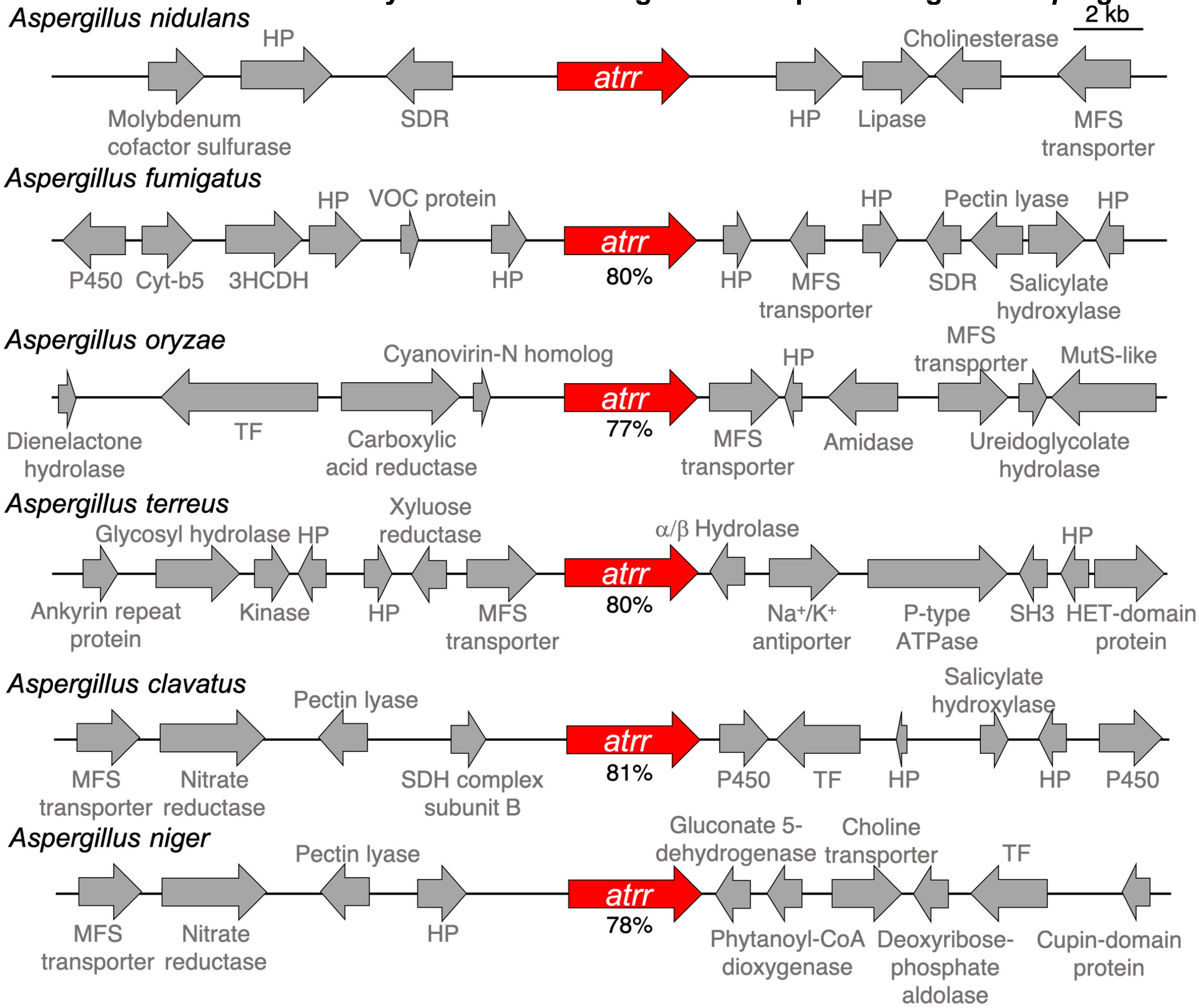

Fig. S3. Comparative genomic context analysis of *atrr* orthologues from closely-related species of the same genus, *Aspergillus*.

### 2.4. Sequence alignment of ATRR-A from different fungal orthologues.

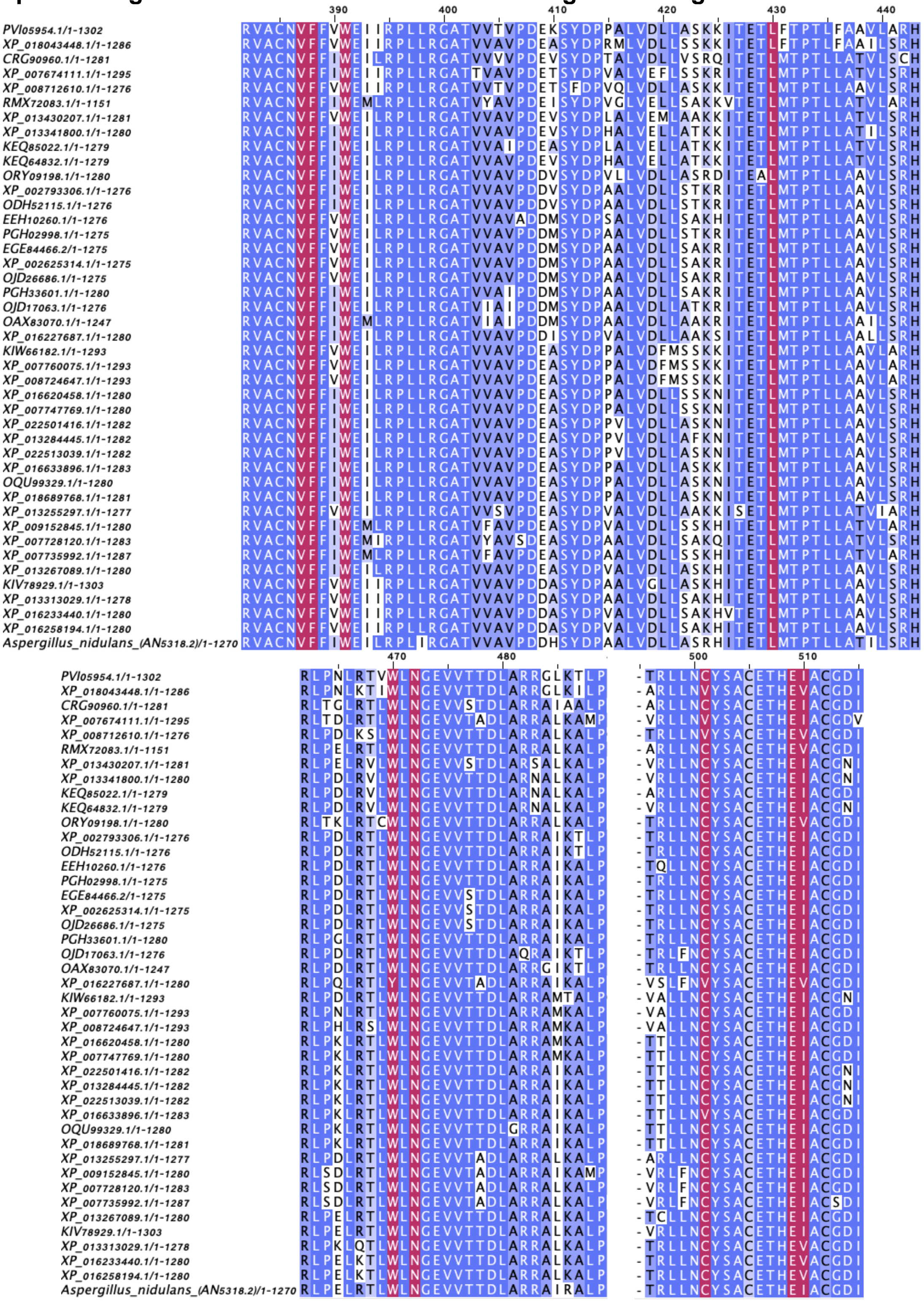

Fig. S4. Continued on next page.

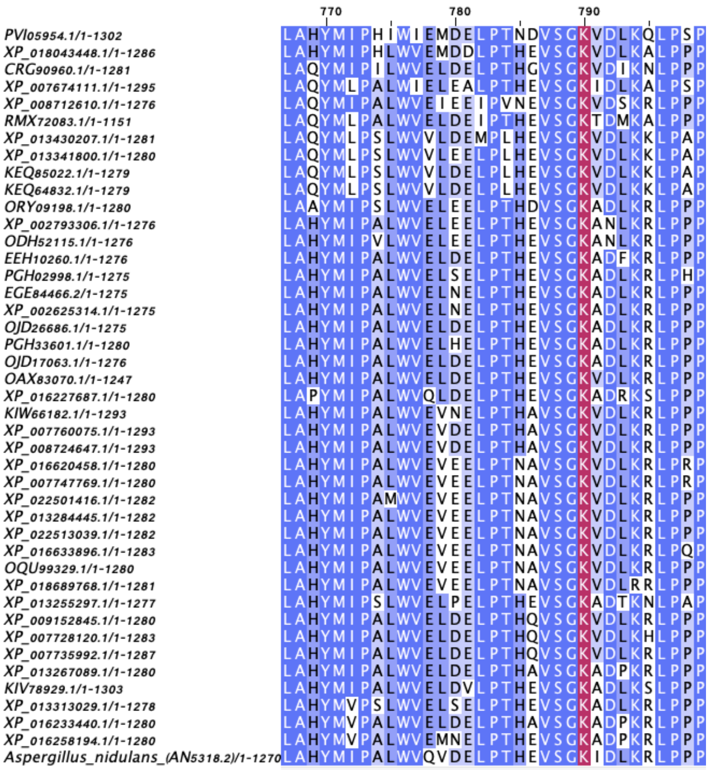

Fig. S4. Multiple sequence alignment of A domain of ATRR orthologues. The accession numbers for selected sequences are shown. The sequences are colored based on sequence identity and the 10AA code residues are highlighted in maroon.

### 2.5. Library of carboxylic acids for in vitro enzymatic activity screening

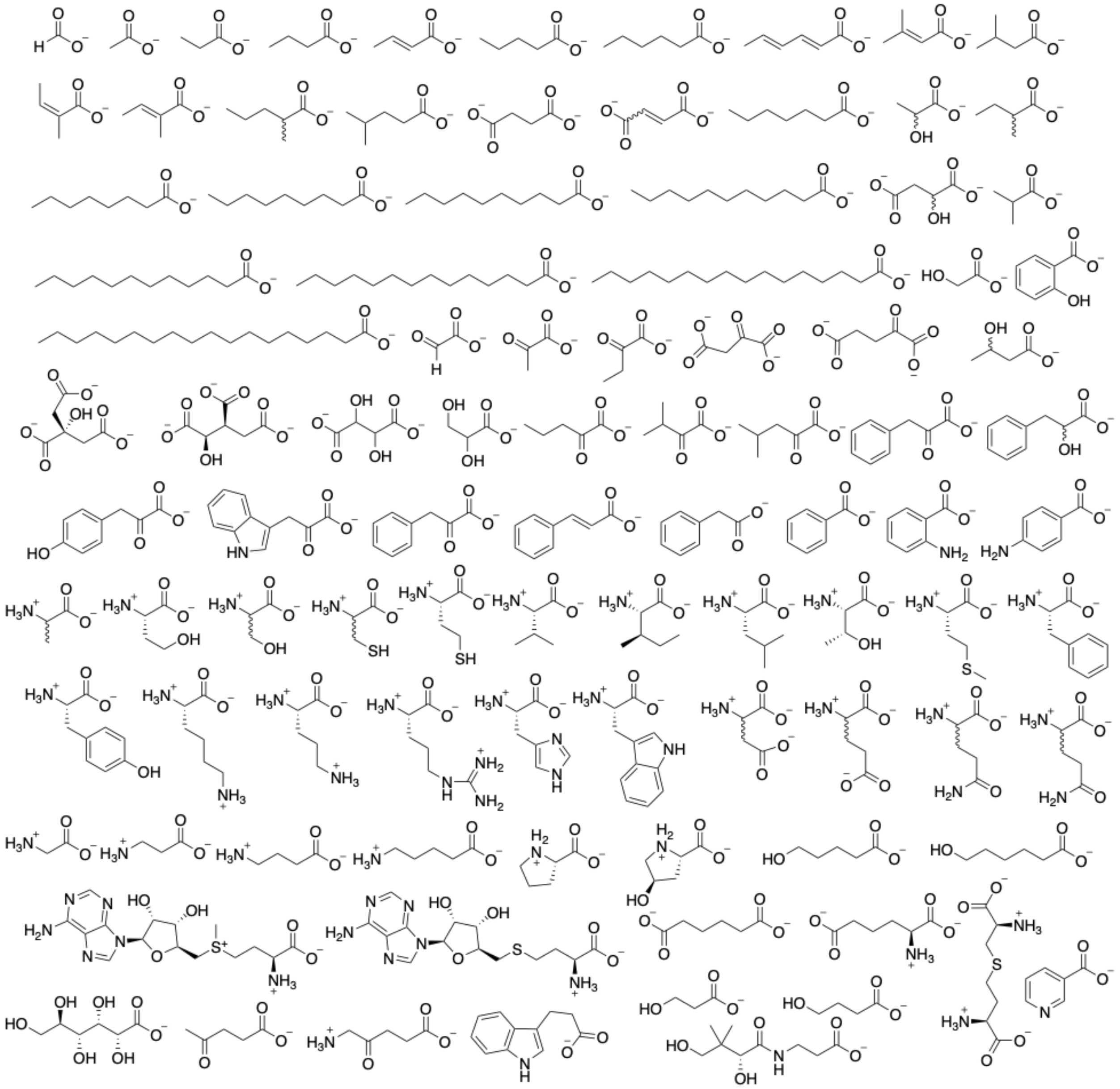

Fig. S5. Chemical structures of the carboxylic acids library used for initial in vitro activity screening.

**2.6. SDS-PAGE analysis of protein purity.**

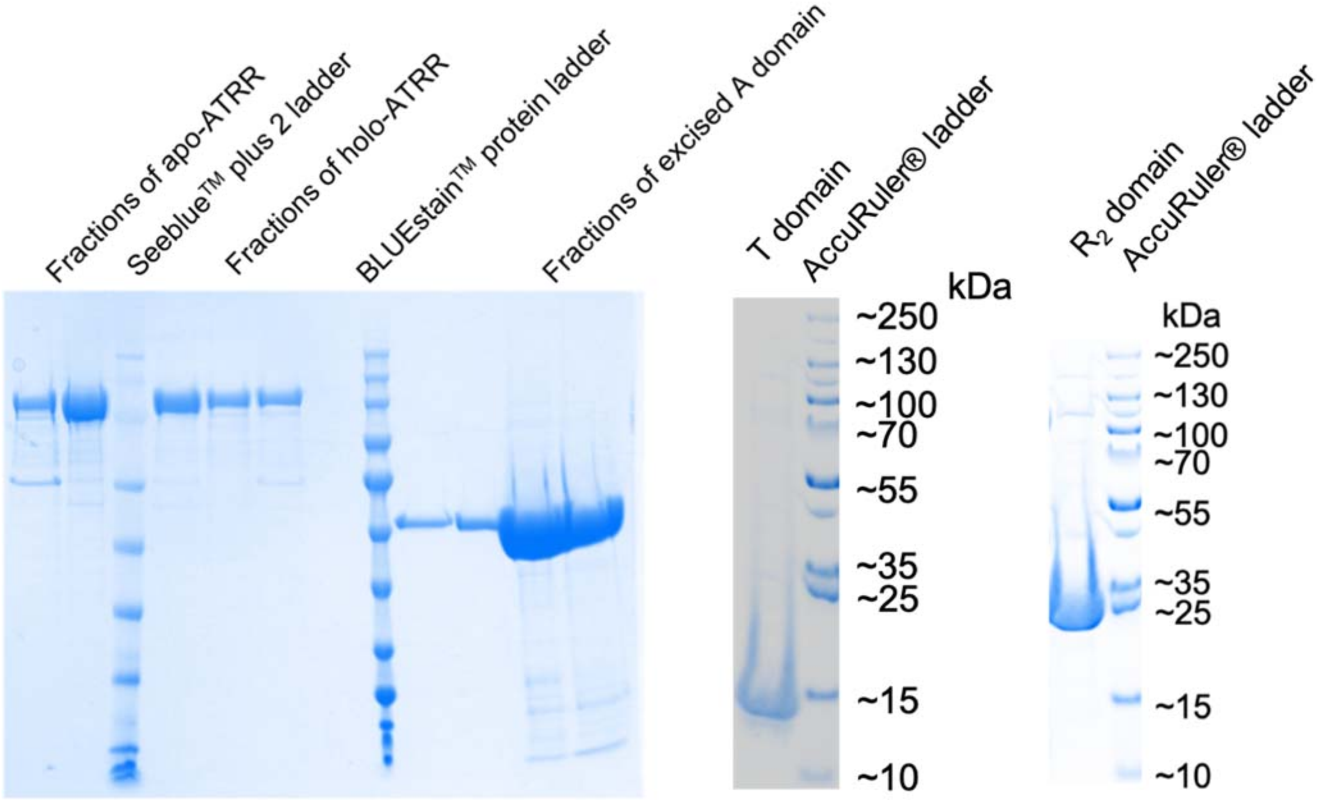

Fig. S6. SDS-PAGE analysis of purified ATRR protein, and dissected domains.

### 2.7. MALDI-TOF characterization of standalone T domain.

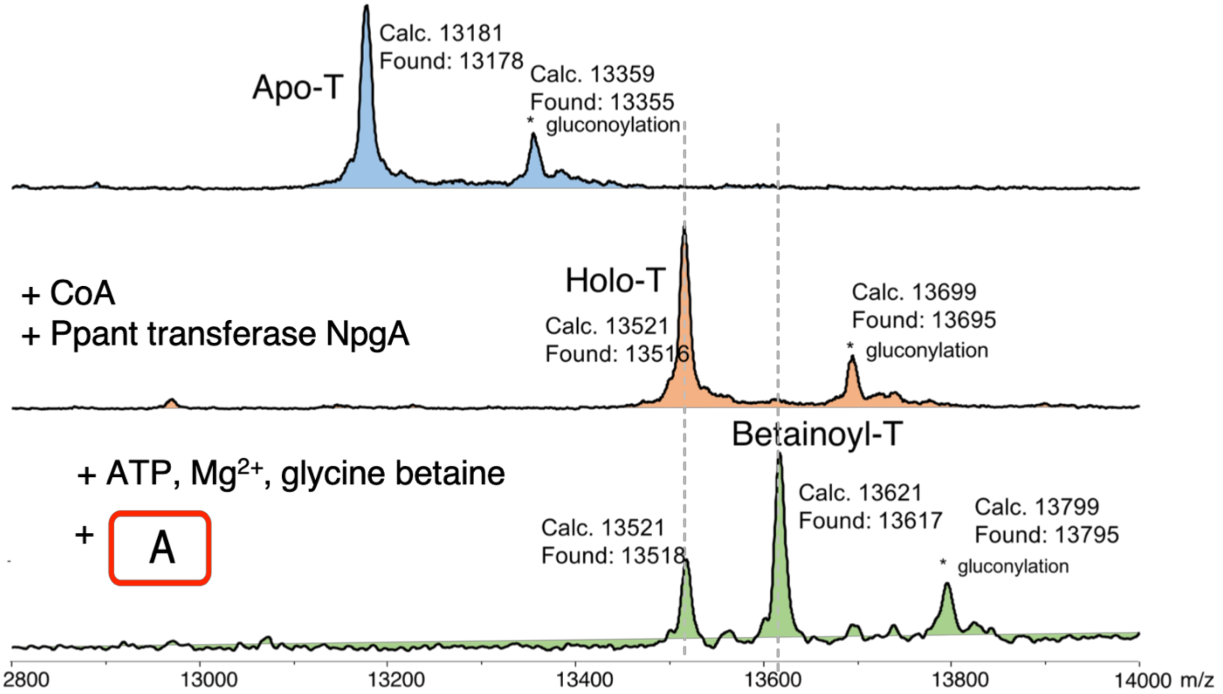

Fig. S7. MALDI-TOF analysis of standalone T domain. Apo-ATRR can be converted to Holo-ATRR enzymatically by NpgA. Holo-T domain can be charged with glycine betaine by standalone A domain.

### 2.8. Choline-derivatization and LC-MS characterization

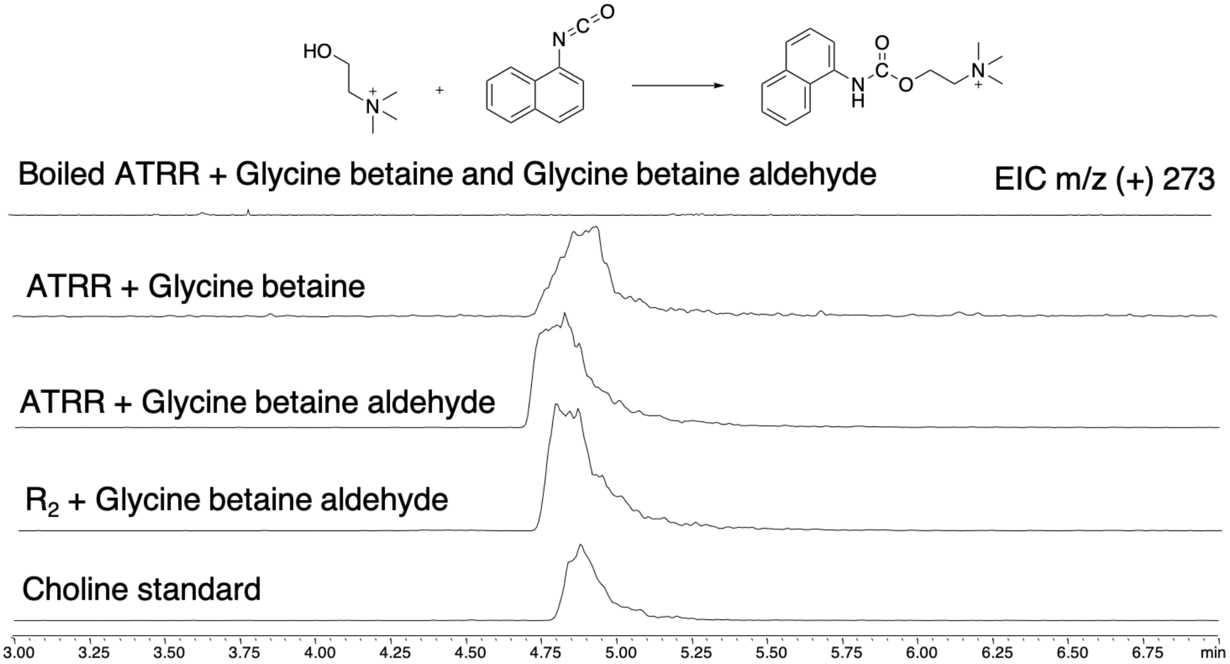

Fig.S8. Choline is accumulated in the assay mixture and the identity was verified by LC-MS after derivatization.

### 2.9. Biochemical characterization of glycine betaine reductase activity

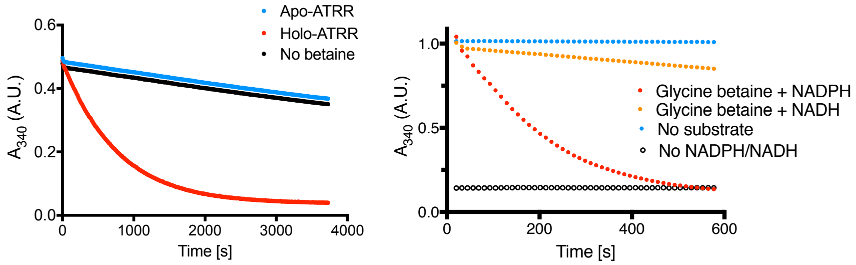

Fig. S9. Biochemical characterization of glycine betaine reductase activity. (left) Phosphopantetheinylation dependency. (right) Cofactor preference. Higher enzyme concentration was required to observe NADH oxidation.

### 2.10. Steady-state kinetics of ATRR

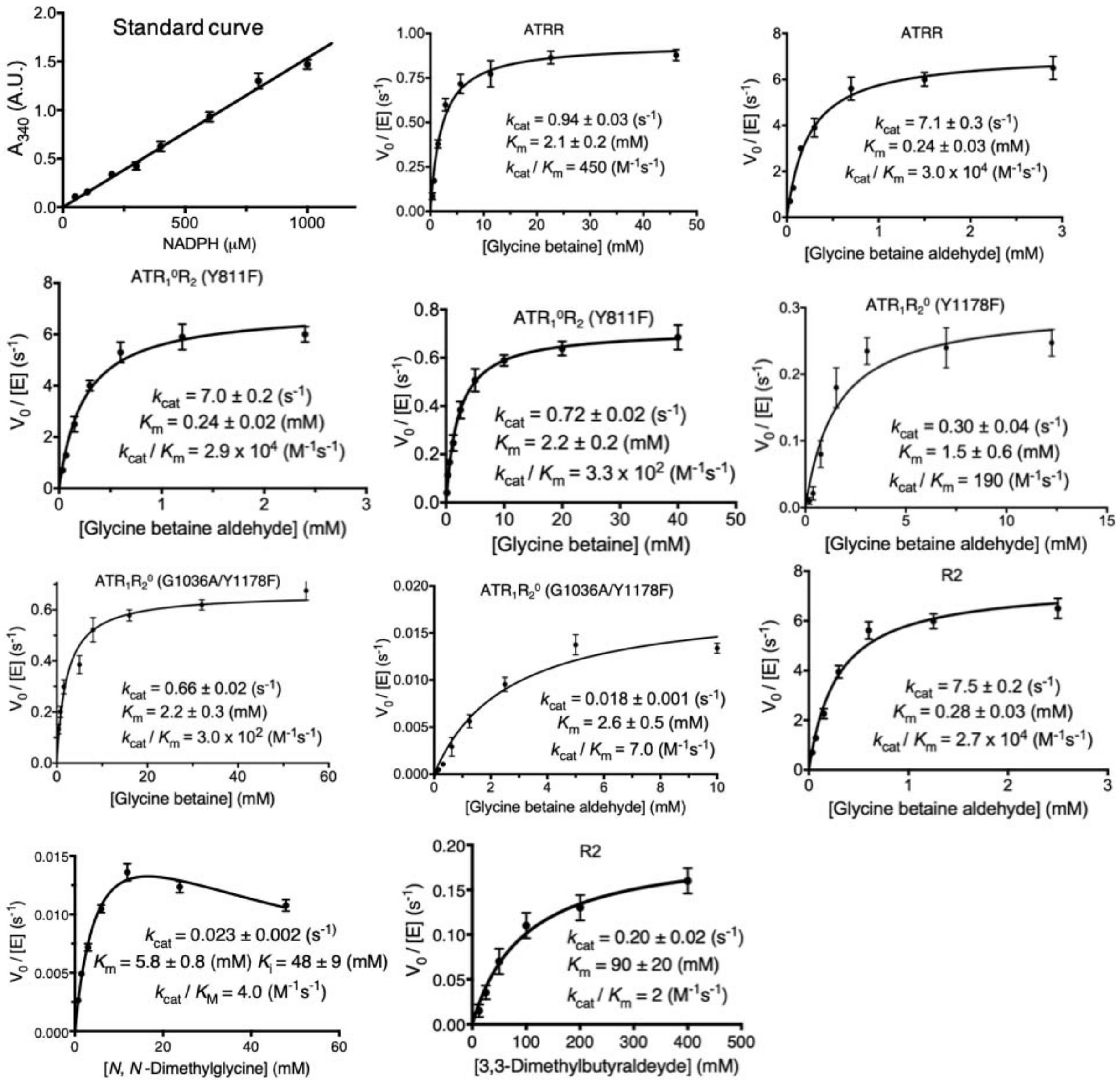

Fig. S10. Steady-state kinetics of ATRR glycine betaine reductase and glycine betaine aldehyde reductase activity. The initial rate was measured based on monitoring the consumption rate of NADPH at 340 nm.

### 2.11. Glycine betaine aldehyde trapped by phenylhydrazine

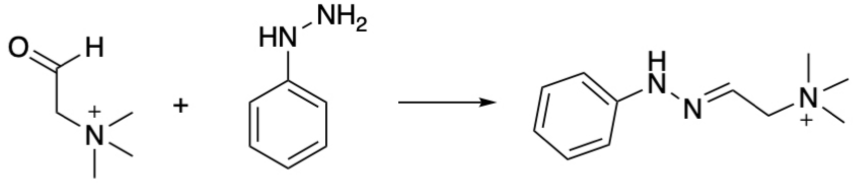

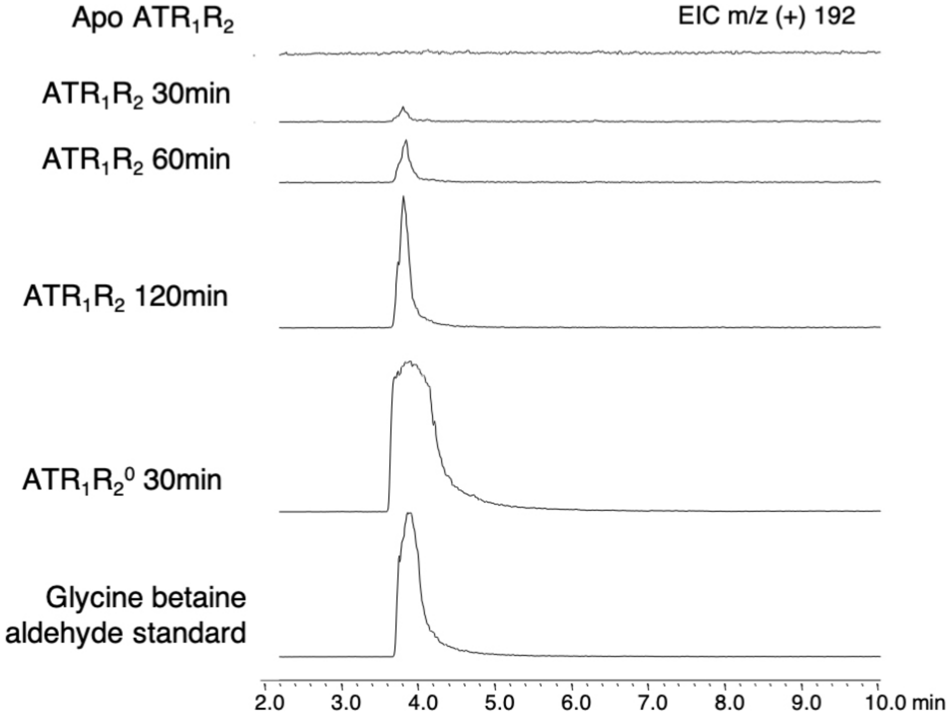

Fig. S11. Glycine betaine aldehyde formation in vitro was verified by LC-MS after derivatization. Phenylhydrazine was included at the beginning of the assay (t=0 min).

### 2.12. Inhibition study of ATRR

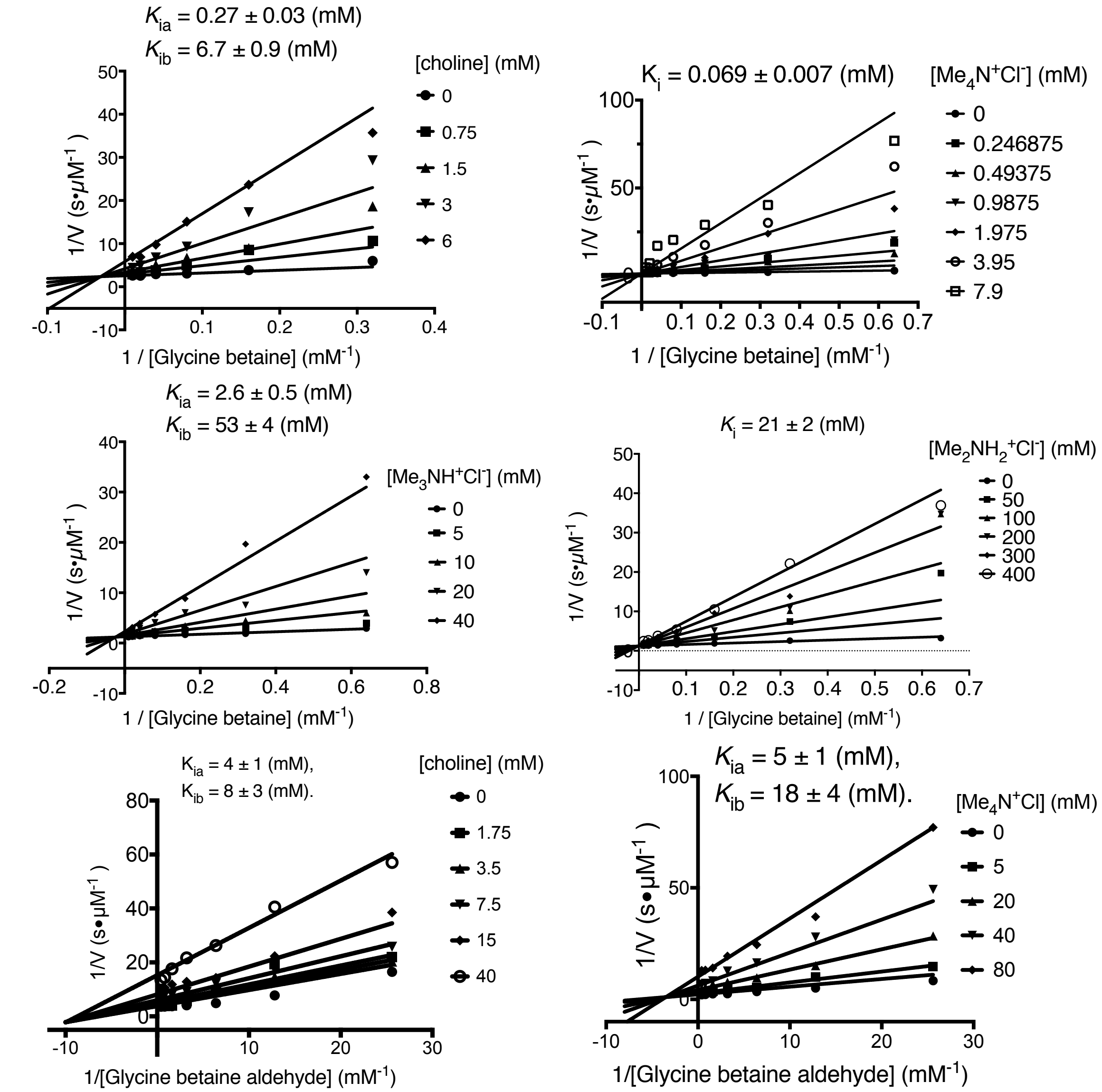

Fig. S12. Lineweaver-Burk plots of ammonium inhibitors of ATRR activity.

### 2.13. Generation of the *atrr* deletion mutant of *A.nidulans*.

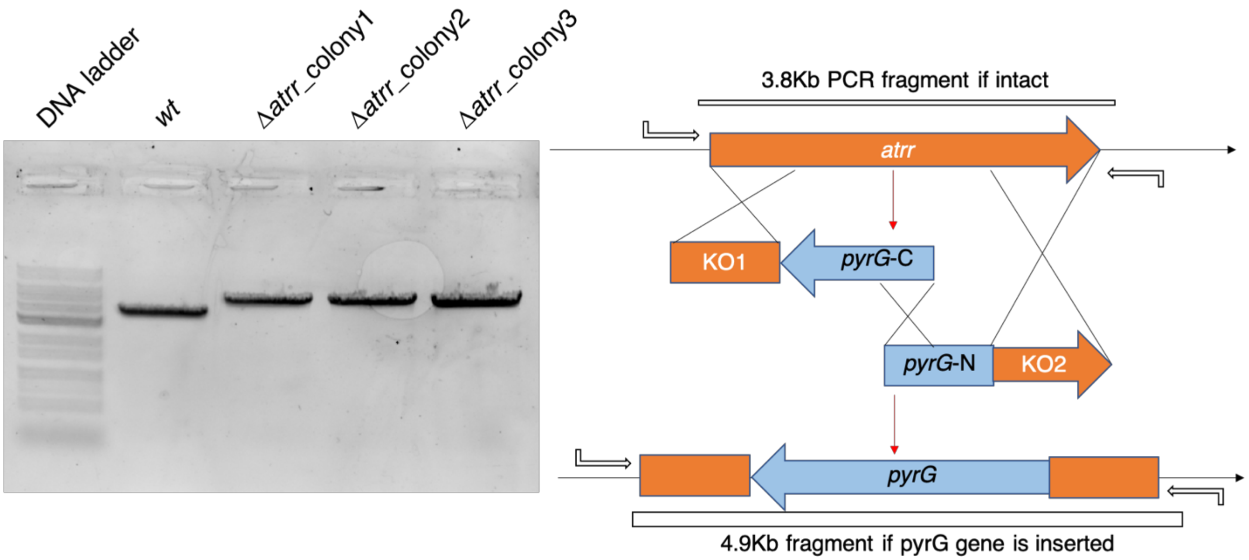

Fig. S13. Deletion of *atrr* gene in wild-type *A. nidulans*. PCR verification (left) and the scheme of split-marker approach (right). Successful gene deletion will cause size change of PCR fragments. Sanger-sequencing was performed on PCR-fragments from confirmed pure transformants.

### 2.14. Generation of the Δ*choA* mutant and Δ*atrr*Δ*choA* double-mutant of *A.nidulans*.

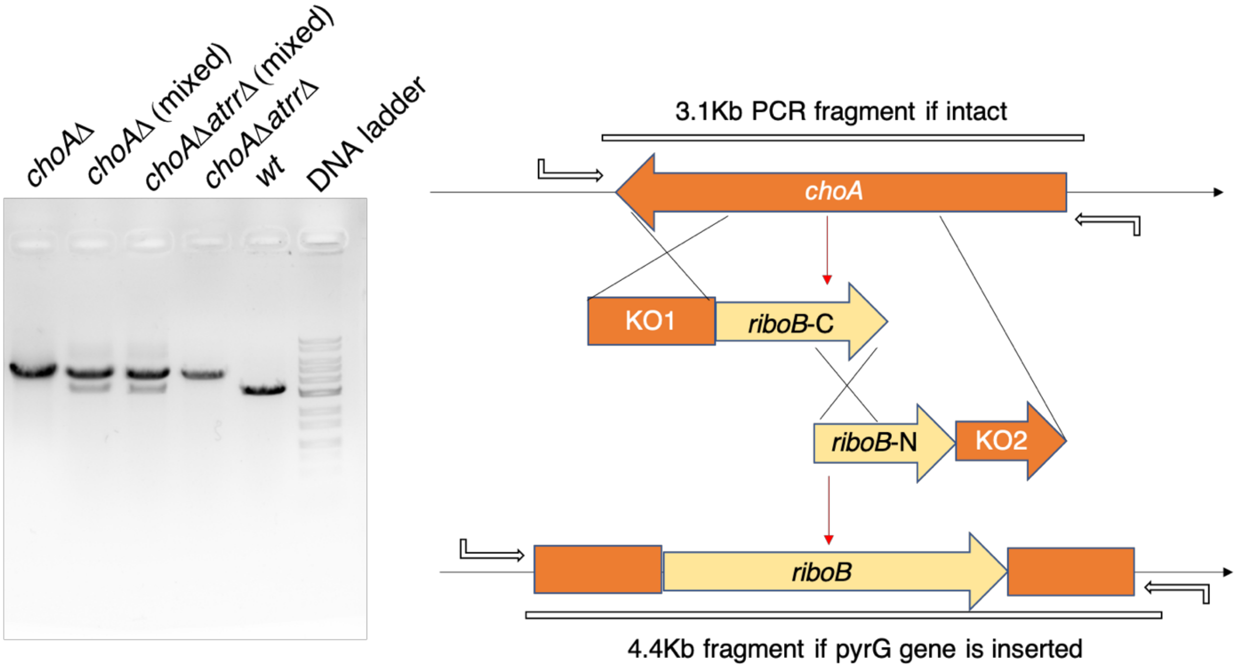

Fig. S14. Deletion of *choA* gene in both wild-type and Δ*atrr* mutant strain of *A*. *nidulans*. PCR verification (left) and the scheme of split-marker approach (right). Successful gene deletion will cause size change of PCR fragments. Successful gene deletion will cause size change of PCR fragments. A double band indicates a mixture of gene deletion and intact loci (mixed transformants). Sanger-sequencing was performed on PCR-fragments from confirmed pure transformants.

### 3. Tables

**Table S1. Oligomers used in this study**

| Name | Sequence |
|---|---|
| ATRR-FL-F | ACCTGTACTTCCAATCCAATGCCATCATTGACACCACAAAAGAC |
| ATRR-FL-R | ATCCGTTATCCACTTCCAATTTAGATTGGCTCGTCTCGCGG |
| ATRR-A-R | ATCCGTTATCCACTTCCAATTTAAGGAAGGCGTTTGAGATCAATC |
| ATRR-T-F | ACCTGTACTTCCAATCCAATGCCAACGGGAACGGAAAGAAGGAAG |
| ATRR-T-R | ATCCGTTATCCACTTCCAATTTATGTGGCATCTGTGCGAAGAACTG |
| ATRR-$R_2$-F | ACCTGTACTTCCAATCCAATGGTCCTCTTAGCGGCAAAGTAG |
| ATRR-G1036A-F | GCAGTCGTTACTGGCGCTTCGTCTGGTATC |
| ATRR-G1036A-R | GATACCAGACGAAGCGGCAGTAACGACTGC |
| ATRR-Y1178F-F | CAGGACTGGGAGTATTTTCGGCCAGTAAGTTC |
| ATRR-Y1178F-R | GAACTTACTGGCCGAAAATACTCCCAGTCCTG |
| ATRR-Y811F-F | GCTGCTGGATGGGTTCGGACAAACTAAGTG |
| ATRR-Y811F-R | CACTTAGTTTGTCCGAACCCATCCAGCAGC |
| *atrr*-KO-F1 | ATGGCCATCATTGACACCACAAAAGACCTT |
| *atrr*-KO-R1 | GCAATTCTGATAACAGATGGTGCCATTGTCTGGGCCCGCCAACGCA GAGTTCACCAG |
| *atrr*-KO-*pyrG*-F | CTGGTGAACTCTGCGTTGGCGGGCCCAGACAATGGCACCATCTGTT ATCAGAATTGC |
| *atrr*-KO-*pyrG*-R | CACCATCGCCACCTTTGACGTGACCCCAGAAAATTACGGGTATGTC CTCCACTGTG |
| *atrr*-KO-F2 | CACAGTGGAGGACATACCCGTAATTTTCTGGGGTCACGTCAAAGGT GGCGATGGTG |
| *atrr*-KO-R2 | CTAGATTGGCTCGTCTCGCGGCTCAATCAGGATCTCG |
| *atrr*-KO-splitR | CAATACCGTCCAGAAGCAATACCACGGCGGTG |
| *atrr*-KO-splitF | CTTTCTGGTACCGCTGTGCAGCTTCAACC |
| *atrr*-KO-checkF | CCAGAATCTGACGTTCTGCTTCCATCGCTAATCG |
| *atrr*-KO-checkR | GCGACATGGACCCAATGCGACAAATGCAAGTC |
| *choA*-KO-F1 | TCACTCTTTGCTTTCAGTTGGTG |
| *choA*-KO-R1 | GTGCGTCATTTATACGACCTACGAGATTACTCGC |
| *choA* -KO-*riboB*-F | GCGAGTAATCTCGTAGGTCGTATAAATGACGCACTGCCACCACG |
| *choA* -KO-*riboB*-R | GCTATAAAGTGCTGTTCATCTGGATGACTCCCTTTCGATGATACCC |
| *choA*-KO-F2 | ATCGAAAGGGAGTCATCCAGATGAACAGCACTTTATAGCTTGC |
| *choA*-KO-R2 | ATGGATCGTGGCCTCTCAACAG |
| *choA*-KO-splitR | GCTCGGATACCCACGACAACTGG |
| *choA*-KO-splitF | CCTTCTTGGCGCAGGTACAC |

| | |
|---|---|
| *choA* -KO-checkF | GGTTGCTTTTCGTGGTTCAGTTTTATCATTCC |
| *choA* -KO-checkR | CTGTTTCATTCTCTAACTTCTTTCTTCTCTTCAACGCTC |
